# Long-term stability of soil microbiome structure and function in liquid soil extracts

**DOI:** 10.64898/2026.05.29.728658

**Authors:** Siqin Li, Genesis Nicole Carpio Paucar, Stokley Voltmer, Natalie J. Kay, Amelia Sadlon, Natalie G. Farny

**Affiliations:** Program in Bioinformatics and Computational Biology, Worcester Polytechnic Institute, Worcester, MA, United States; Department of Biology and Biotechnology, Worcester Polytechnic Institute, Worcester MA, United States

**Keywords:** Soil extracted soluble organic matter (SESOM), soil microbial community (SMC), network analysis, synthetic biology

## Abstract

Soil microbial communities (SMCs) play an important role in various ecological processes, including plant growth, carbon cycling, and greenhouse gas production and consumption. There have been many prior studies of soil microbiome function and structure. However, soil is a complex environment in which to conduct biological studies. Therefore, simplified SMC models, often adapted to liquid culture, have been employed in the laboratory to study specific microbial interactions and individual microbial functions. Specific advantages of these laboratory liquid SMC models include the ability to modulate community membership, control environmental conditions, and employ high-throughput assay techniques. The disadvantages of current laboratory liquid SMC models include long cycles for growing bacteria *in vitro*, the obligatory use of strains that are culturable in isolation, intricate media requirements, and complex community assembly protocols. To address some limitations of current liquid SMC models, we sought to create a streamlined process for extracting and maintaining a liquid culture of an existing SMC. Soil-Extracted Solubilized Organic Matter (SESOM) was made from four different soil types, including rich organic potting soils and environmental samples, and filtered to maintain the SMC. These SESOM liquid SMC models were cultured for 28 days, and SMC composition was measured by 16S rDNA sequencing. The SESOM SMCs maintain high alpha and beta diversity over time, including strains that are not culturable in isolation, with the greatest stability correlated with higher soil organic carbon. Further, the SESOM SMCs maintain unique signatures of their starting solid soils, suggesting that drift in SMC composition over extended time in liquid culture does not eliminate the defining microbial relationships of a given soil type. Network analysis of SESOM SMCs relative to solid soils suggests the functional roles of bacterial taxa were maintained in the liquid models over time. We further demonstrate that the platform can be applied to monitor the survival and persistence of a model engineered microbe – the common synthetic biology chassis *Pseudomonas putida* – within a native SMC. We conclude that the SESOM model is a valuable tool for facilitating the study of SMCs in the laboratory.

## INTRODUCTION

Soils harbor some of the most diverse microbial communities on Earth. These diverse soil microbial communities (SMCs) are involved in essential ecosystem processes that maintain soil health. The complexity, organizational structure, and diverse functions of SMCs across the globe are gradually being revealed (Delgado-Baquerizo et al., 2018, 2020; Egidi et al., 2019; Labouyrie et al., 2023). It has become increasingly clear that SMCs play important roles in a broad range of processes including nutrient cycling, soil structural integrity and stability, promoting plant growth, participating in biocontrol of pathogens, bioremediation, and greenhouse gas regulation (Delgado-Baquerizo et al., 2025; Iqbal et al., 2025; Philippot et al., 2024a).

While our knowledge of SMCs has grown, so too has grown the scope and intensity of threats to soil health. An estimated 33% of global soils are in a state of moderate to severe degradation (Smith et al., 2024). Soil degradation is the result of a variety of factors including erosion, unsustainable management practices, pollution, salinization, and more (Timmis & Ramos, 2021). Synthetic biology offers the promise of biotechnologies to advance soil health. A particularly notable example is the genetic engineering of a field strain of *Klebsiella variicola* to enhance nitrogen fixation (Wen et al., 2021). This genome edited strain has been applied to millions of acres of U.S. farmland to reduce the need for synthetic fertilizer, enabling a significant decrease in synthetic fertilizer application with equivalent or even improved crop yields (Woodward et al., 2025). Similar biotechnology-based approaches are needed, in conjunction with changes in management practices, to address the broad host of growing threats to global soil health.

One barrier to the rapid development of new microbial biotechnologies for soil applications is the lack of flexible and scalable laboratory models of SMCs (Zhalnina et al., 2018). Most laboratory models to date have taken the approach of re-building SMCs from culturable members (known as synthetic communities, or SynComs) using defined soil-like medium. Using a simplified seven-member maize root SynCom, the impact of the SMC on fungal disease resistance in maize was demonstrated (Niu et al., 2017). Similarly, SynComs of sixteen to seventeen members have been assembled and tested in *in vitro* plant-microbe models, and have been shown to have great reproducibility (Coker et al., 2022; Novak et al., 2025). However, while these SynCom models have contributed enormously to our understanding of SMCs, particularly in the context of rhizobial relationships, they suffer from some specific limitations as a platform for synthetic biology. These limitations include time-consuming assemblies, limitation to only microbial members that can be cultured in isolation, and low-throughput processes. To enable rapid prototyping of engineered bacterial members and assess their survival and function within existing SMCs, additional approaches are needed that retain the characteristics of native SMCs while enabling high-throughput measurement and rapid iteration.

A promising approach to bridge this gap in laboratory models of SMCs is the use of liquid soil extracts in the form of Soil Extracted Solubilized Organic Matter (SESOM). SESOM has been used as a sterile growth medium for soil microbes in both liquid cultures and plates (Brözel et al., 2011; Luo et al., 2007; Vilain et al., 2006a), and can be made from any soil. The chemical composition of SESOM depends upon the soil physicochemical characteristics, which can be characterized, and correlates with the initial soil properties (Liebeke et al., 2009). The SESOM model has been used for a variety of applications, from the study of rhizobial bacterial survival and dynamics (Sandhu et al., 2021, 2023, 2025; White et al., 2015) to the persistence of pathogenic *E. coli* in soil (NandaKafle et al., 2017, 2018). Liquid soil models are of particular value to the synthetic biology design-build-test-learn (DBTL) cycle, as they enable high-throughput approaches such as automated liquid handling and flow cytometry.

Here, we describe a non-sterile preparation of SESOM wherein we have isolated the microbiomes from both rich commercial soils and environmental soils. These liquid-extracted SMCs have high similarity and diversity relative to the starting soils, which is maintained over a 28-day culture period. We apply network analyses to demonstrate the overall taxonomic stability of liquid SMCs relative to their starting soils over time, as well as conservation of functional pathways within communities. Finally, we demonstrate that the survival and persistence of an engineered microbe – the environmental synthetic biology chassis organism *Pseudomonas putida*, expressing the fluorescent protein tdTomato – can be monitored over extended time in co-culture with a native SMC using SESOM.

## MATERIALS AND METHODS

### Soil samples

Four different types of soils were used, categorized to two groups. Samples from commercial sources were selected based on their “organic” designation (BB, black bag sample, Miracle-Gro Performance Organics Container Mix, and GB, green bag sample, Espoma Organic Potting Mix, Supplementary Fig. S1A-B). Environmental samples were taken at a private property in Massachusetts (approximately 42.264835337557585, -71.35118446038254). The area is a suburban neighborhood. Sample 1 was taken from an area at the corner of the driveway to the property, immediately bordering the street (roadside sample, RS, Supplementary Fig 1C). Sample 2 was taken from a wooded area between two properties that has remained undisturbed for at least 15 years and is approximately 100 yards from the roadside sample (wooded sample, WS, Supplementary Fig. 1D). Leaf litter was cleared, and soil from a hole ∼8 inches deep by 6 inches wide was collected.

### Soil analysis

Soil samples were sent to the Soil and Plant Nutrient Testing Laboratory at University of Massachusetts Amherst for testing. The analysis included soil organic matter, soluble salts, nitrate and scoop density, pH, cation exchange capacity and nutrients. The basic physicochemical properties of each soil are shown in Supplementary Table S1.

### SESOM (Soil-Extracted Solubilized Organic Matter)

Soil extracted solubilized organic matter (SESOM) was prepared as described (Vilain et al., 2006, Fig. 1), with the following modifications. 100g of soil was prepared in 500mL of 1xPBS in a 1L flask, and placed on a shaker at 200 RPM, 37° C for 2 hours. Large particulates were separated using a french press. The remaining mixture was filtered under vacuum in a Buchner funnel through two layers of Whatman paper, grade 1 and grade 4, to create a particle retention size of approximately 11 µm. This preparation is known as non-sterile SESOM (NS-SESOM). To generate sterile SESOM (ST-SESOM), the 11 µm filtrate was filtered through an 0.22 µm PES vacuum filtration system.

**Figure 1.**
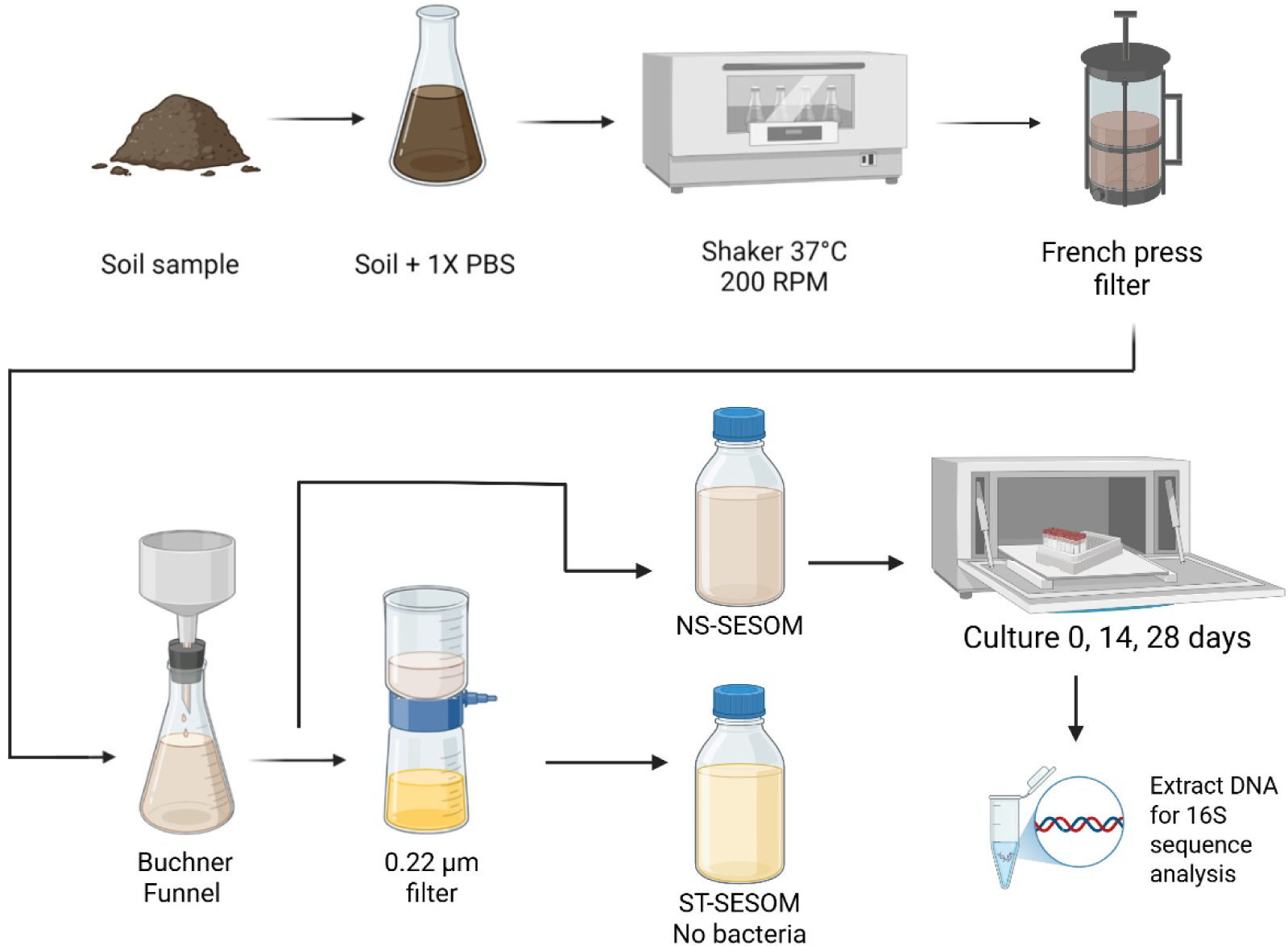
Overview of SESOM generation and sample culture and sequencing protocol. See Methods section for details. Figure generated with Biorender.

### DNA extraction

To generate DNA samples for sequencing, 25 mL of non-sterile SESOM was centrifuged at 32,000 rpm for 30 minutes at 25°C, then DNA was extracted using the Qiagen PowerSoil Pro DNA extraction kit per the manufacturer’s instructions.

### 16S rDNA sequencing and processing

The sequencing fastq files were generated by 2×250 Illumina MiSeq amplicon sequencing of the V3-V4 hypervariable regions of the 16S rDNA by Azenta Life Sciences.

### Taxonomic Identification

The DADA2 pipeline (Callahan et al., 2016) with QIIME2 software (Bolyen et al., 2019) was used to assign amplicon sequencing variants (ASVs). Due to the sequence error at the end of reads, forward reads were retained as original and reverse reads were truncated to 230bp. Since primer sequences are a part of 16S rDNA sequences, they are kept for the analysis. Ambiguous bases (N) were discarded from the sequences. Reads with more than 2 expected errors were also removed. Bases with a quality score (Q) below 2 were eliminated. The error model was estimated based on filtered forward and reverse reads and used for denoising process. Then read pairs were merged based a minimum overlap of 20 bp and merged reads with more than 2 mismatches in the overlapping region were removed. The chimera sequences were also discarded from sequences. Then the dataset was cleaned out for downstream analysis. The soil samples were annotated with SILVA 138 reference database (Quast et al., 2013). Amplicon sequencing variants (ASV) were assigned using a pre-trained classifier SILVA 138, which is a taxonomic classifier trained on the SILVA 138 database using naïve Bayes method. The soil taxonomy composition was the average relative abundance of soil microbial community for all duplicates under same condition. Taxonomy results were generated by QIIME2. The taxa bar plots were visualized with ggplot2 package (Wickham, 2011).

### Diversity analyses

Shannon index to describe alpha diversity (Kim et al., 2017) and weighted UniFrac to describe beta diversity (Anderson et al., 2006) were visualized with Principal Coordinates Analysis (PCoA) plot by ggplot2 package (Wickham, 2011).

### Network analysis

Co-occurrence networks were built for each soil type sample at four different conditions, including solid soil at day 0 and NS-SESOM at day 0, day 14 and day 28. Networks were constructed on the class level. Bacteria taxa were represented by nodes and edges between nodes represented the interactions between two different taxa. Positive or negative relationships were calculated by Pearson correlation with centered log-ratio transformation normalization, further accounting for compositionality. Zero values were handled with multiplicative replacement. Networks were constructed based on the 50 taxa with highest frequency at the class level. Network construction and analysis was performed and visualized using the NetCoMi package (Peschel et al., 2021).

To compare the similarity between networks, global network properties were calculated. These included the clustering coefficient, modularity, positive edge percentage, edge density, and natural connectivity. The comparison of network also included network nodes. The network nodes were compared across different conditions in each soil type and the results were visualized with Venn diagram by ggvenn package (Gao et al., 2024). Node colors represented the determined clusters and node sizes were scaled according to the nodes’ eigenvector centrality. Hubs are nodes with an eigenvector centrality value above the empirical 95% quantile of all eigenvector centralities in the network. Red edges represented negative correlations and green edges represented positive correlations between bacteria in the communities.

### Metagenome Functional Predictions

The functional profile of the SMC was predicted by PICRUSt2 (Phylogenetic study of Communities by Reconstruction of Unobserved States) (Douglas et al., 2020). Then, the network was built with NetCoMi based on the inference of functional pathway abundance, as described above. Each node represents a specific pathway, and edge represents the correlations between pathways. The same parameters were chosen to build the functional network as for taxonomic networks.

### Flow Cytometry

*Pseudomonas putida* AG4775 was a gift from Dr. Adam Guss, Oak Ridge National Laboratory. AG4775 is a derivative of KT2440 containing a multi-site recombinase system (Elmore et al., 2017). tdTomato was stably integrated into the genome as a fluorescent marker using the Bxb recombinase as described (Elmore et al., 2017).

Sterile or non-sterile SESOM preparations were generated as described above. *P. putida* with an integrated tdTomato marker was grown in an overnight culture of LB at 30°C shaking at 220 rpm, then pelleted and resuspended in sterile 1x PBS for measurement of optical density (OD₆₀₀). Cells were diluted into 3 mL cultures of NS-SESOM or ST-SESOM at an OD_600_ = 0.5, and cultured in a 24-well deep well plate at 30°C shaking at 220 rpm. Samples were removed and diluted 1:40 in 1xPBS containing 200 µg /mL kanamycin. Each 200 µL dilution was dispensed into individual wells of a flat-bottom 96-well plate, and measured by flow cytometry at the timepoints indicated on a Beckman Coulter CytoFlex flow cytometer using the following settings: FSC (Forward Scatter): 500; SSC (Side Scatter): 1000; RFP (Red Fluorescent Protein): 100; flow rate to 20 µL/min, 20 µL sample volume. Analysis was performed using FlowJo software (FlowJo LLC). Flow cytometry results were compiled using GraphPad Prism v10.6.1.

### Colony Forming Unit (CFU) Assays

10g of soil was weighed into a glass petri dish. Sterile soils were wrapped in aluminum foil and autoclaved at 121° C for 30 minutes. An overnight culture of *P. putida* expressing tdTomato and a kanamycin resistance marker was centrifuged to remove LB medium, then resuspended in 1X PBS at a density of OD_600_ = 0.1 per mL. 1.0 OD_600_ of cells (10 mL at OD_600_ = 0.1 per mL) of was added to the 10 g of soil in each plate and mixed well with a metal spatula. For CFU sampling, an approximate 0.5 g soil sample was removed and weighed. 1X PBS was added at a volume of 5 mL per 0.5 g, scaled accordingly for the exact weight of the sample. Samples were vortexed vigorously, then filtered using a 5-micron syringe filter to remove large soil particulates. Samples were then serially diluted and 100 µL of dilutions were plated to LB agar with 50 µg/mL kanamycin, incubated overnight at 30° C, and then colonies were counted and recorded. Counts were used to calculate colony forming units (CFU) per g of soil.

### Data processing and statistical analysis

Linear mixed-effects models (LMMs) were used to model the longitudinal soil microbiome dataset. LMMs was modeled using QIIME2 to determine the relationship between taxa abundance or diversity index (alpha or beta diversity). The association was built between the response variable (top 5 most abundant taxa, alpha diversity, or beta diversity) and soil community trait variable (indicating either liquid SESOM or solid soil model, or time), as indicated for each analysis.

All code used to process and analyze sequencing results and perform the data visualization can be accessed through Github at https://github.com/FarnyLabWPI/soil-metagenomics.

Flow cytometry and CFU assays were analyzed and visualized using GraphPad Prism v10.6.1.

## RESULTS

### Non-sterile SESOM (NS-SESOM) extracts maintain high fidelity and microbial diversity over time in culture

Many prior studies have reported the use of a sterile preparation of SESOM as a growth medium to study the growth, survival, and other phenotypes of soil bacterial species (Brözel et al., 2011; Luo et al., 2007; NandaKafle et al., 2017, 2018; Sandhu et al., 2021, 2025). We predicted that it may be possible to maintain in-tact microbial communities in a liquid culture by applying selective filtration to maintain the bacterial community in the preparation of SESOM, which we refer to as non-sterile SESOM (NS-SESOM). 3-(N-morpholino)propanesulfonic acid (MOPS) was used in prior descriptions of SESOM preparation (Brözel et al., 2011; Vilain et al., 2006b; White et al., 2015). However, MOPS is a complex carbon-containing molecule. We observed that our strain of *Pseudomonas putida* KT2440 was able to produce fluorescent protein in MOPS only buffer but was unable to do so in phosphate-buffered saline (PBS) (Supplementary Fig. S2), suggesting *P. putida* may be able to catabolize MOPS. The observation is consistent with reports of environmental microbes with the capacity for morpholine catabolism (Knapp et al., 1996; Poupin et al., 1998), and the known complex catabolic capabilities of *P. putida* (Jiménez et al., 2002; Loeschcke & Thies, 2015; Shrestha et al., 2023). Therefore, 1xPBS was used as the extraction buffer so that the only carbon and nitrogen sources available in the medium would be from the soil extract.

To determine whether NS-SESOM-extracted SMCs were an accurate representation of solid soil SMCs, and to measure the stability of SN-SESOM SMCs over extended time in culture, we performed 16S sequence analysis on solid soils and corresponding NS-SESOM at days zero, 14 and 28 post-extraction (Fig. 1). The analysis was performed with a total of four soils: two commercially available organic potting soil mixes (Black Bag (BB, Fig. 2A) and Green Bag (GB, Fig. 2B)), and two environmental samples taken from a roadside (RS) or a wooded (WS) area in a suburban neighborhood (Fig. 2C-D). The relative abundance of various taxa by phylum (Supplementary Figure S3), class (Fig. 2), and genus (Supplementary Figure S4) are shown. The full taxonomic analysis is available in Supplementary Tables S2-S7.

**Figure 2.**
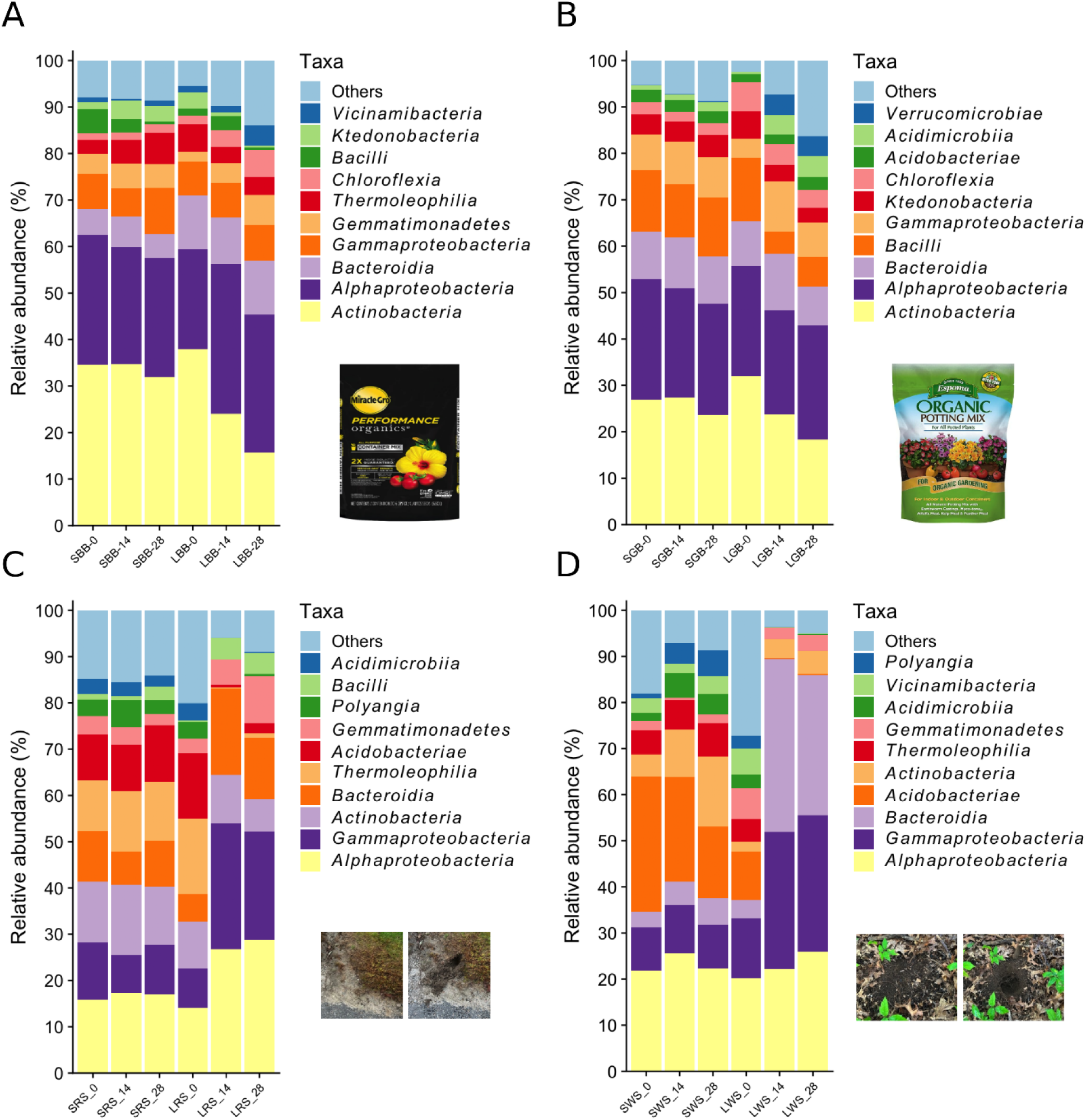
Bacterial abundance by class in soil and NS-SESOM samples. Sample ID tags are assembled as follows: L (liquid) represents NS-SESOM and S represents solid soil samples; commercial samples are BB (black bag, **A**) and GB (green bag, **B**), environmental samples are RS (roadside, **C**) and WS (wooded, **D**). The DNA samples were collected at day 0, day 14 and day 28, as indicated by the number following each sample ID tag, and subjected to 16S sequencing and ASV annotations by class. The 10 most frequent bacteria classes are indicated by colors. Each bar chart represents the average of three sample replicates for each condition.

At all taxonomic levels, the relative abundance of the dominant taxa appeared remarkably stable across solid and NS-SESOM samples at all timepoints in carbon-rich commercial soils (Fig. 2A-B). Further, for all sample types, the initial (day 0) liquid-extracted NS-SESOM SMC is reliably representative of the initial solid soil SMC. The NS-SESOM models sequenced at days 14 and 28 displayed some shifts in the relative abundance of SMC taxa, which were more pronounced in the environmental samples which had poorer organic content (Fig. 2 C-D), but still maintained the core bacterial members in the community as the initial community at day 0. Interestingly, when examined at the genus level, several uncultured taxa were identified among the most abundant genera, some of which were maintained over the 28-day culture period (Supplementary Figure S4). To quantitatively assess the stability of dominant taxa in NS-SESOM over time, we performed linear mixed effects (LME) modeling on the top five ASVs at the class level within each of the four soil samples (Supplementary Fig. S5-S8, Supplementary Table S8). LME modeling revealed that ASV variation was much more generally associated with the timepoint sampled than the sample type group (liquid NS-SESOM versus solid soil), indicating that the abundance of dominant SMC members do change over time in NS-SESOM culture, though are generally not distinct from the abundance in the starting soil, particularly in the rich organic commercial soils. These analyses suggest that the NS-SESOM extracts maintain the fidelity of the starting soil SMC both at the point of initial extraction and, to some extent, the dominance of major taxa over time in culture.

We next compared the alpha diversity of NS-SESOM and solid soil SMCs (Fig. 3). The Shannon index is a commonly used alpha diversity metric used to quantify microbial diversity in a community, that considers both richness (number of species) and evenness (distribution of species abundance) (Shannon, 1948). In all instances, the solid soils showed little change in alpha diversity over the 28-day period. For NS-SESOM made from commercial samples, the alpha diversity was similarly maintained over time. The environmental NS-SESOM samples showed a notable decrease in alpha diversity after 14 to 28 days in culture. However, in those instances, the overall diversity remained very high, higher than in the commercial samples, even despite the observed decrease over time (Fig. 3). LME models revealed no significant effects of sample type group (NS-SESOM versus solid soil), time (0, 14, 28 days) or their interaction on the BB soil Shannon index, but significant time-by-group interactions were observed in Shannon indices in GB, RS and WS soils (Supplementary Fig. S9 and Supplementary Table S9). In all instances, where changes in alpha diversity were indicated by LME models, the overall alpha diversity of samples remained high. These results suggest that diverse native bacterial SMCs can be efficiently extracted and diversity maintained for extended time in culture using the NS-SESOM method.

**Figure 3.**
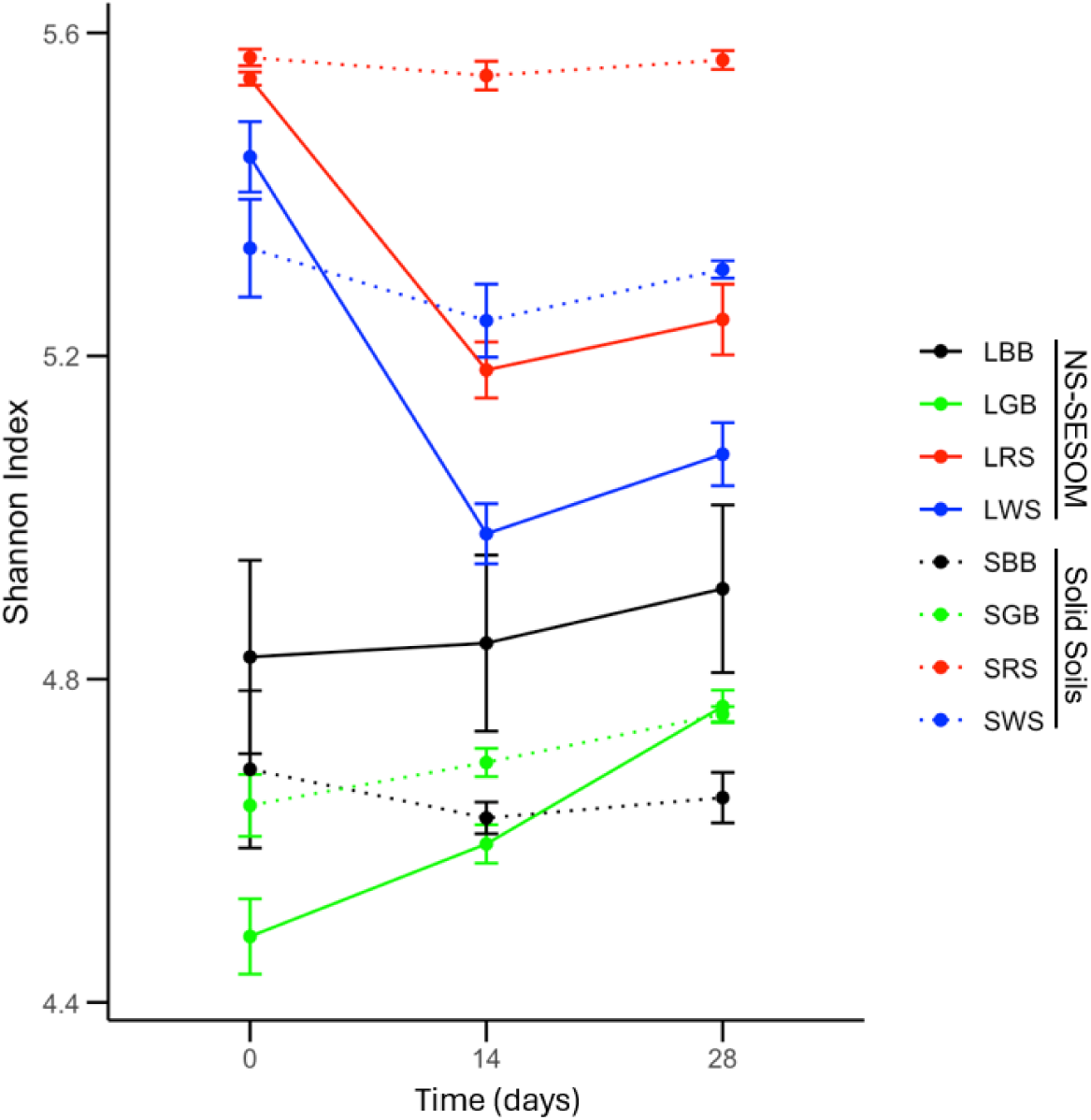
Alpha diversity indices of soils and NS-SESOM samples. Sample ID tags are assembled as follows: L (liquid) represents NS-SESOM (solid lines), and S represents solid soil samples (dashed lines); commercial samples are BB (black bag) and GB (green bag), environmental samples are RS (roadside) and WS (wooded sample). The DNA samples were collected at day 0, day 14 and day 28, and subjected to 16S sequencing and ASV annotations. Shannon index was then calculated for each sample. Error bars represent +/- SEM (n=3).

### NS-SESOM SMCs are reproducible and maintain distinct microbial signatures over time in culture

To determine how the diversity of SMCs compared among different soil types, we performed beta diversity analysis. Weighted UniFrac distance is a commonly used beta diversity metric to assess the relative difference in SMCs between communities, which includes dissimilarity between species abundance and phylogenetic relationships, and were visualized using principal coordinates analysis (PCoA). Beta diversity was first measured within individual soil types, in order to examine the relationship between the NS-SESOM samples and the initial soil SMCs over time. The first two principal component axes (PC1 and PC2) described the majority (69% or greater) of the variation for all soil and NS-SESOM samples within a given commercial or environmental soil type (Fig. 4 A-D). Replicates were clustered under all conditions, indicating the strong reproducibility of the extracted communities in each case. Notably, as observed in the taxonomic analysis (Fig. 2), the solid soil samples separated along PC1 (GB, RS and WS, Fig. 4B-D) or PC2 (BB, Fig. 4A) with the day 0 NS-SESOM samples for all soil types, indicating they shared strong similarity in microbiome compositions based on phylogenetic structure and relative abundance. This result further supports the observation that the initial NS-SESOM preparation is a high-fidelity representation of the starting soil SMC, regardless of soil type. In all soil types, replicates for NS-SESOM at days 14 and 28 clustered away from the solid soil and NS-SESOM at day zero, indicating as anticipated that increasing time in liquid culture is the primary factor impacting SMC composition in NS-SESOM. Consistent with the PCoA results, LME models show that when differences exist, they are more often an effect of sample time rather than sample type (Supplementary Fig. S10 and Supplementary Table S10).

**Figure 4.**
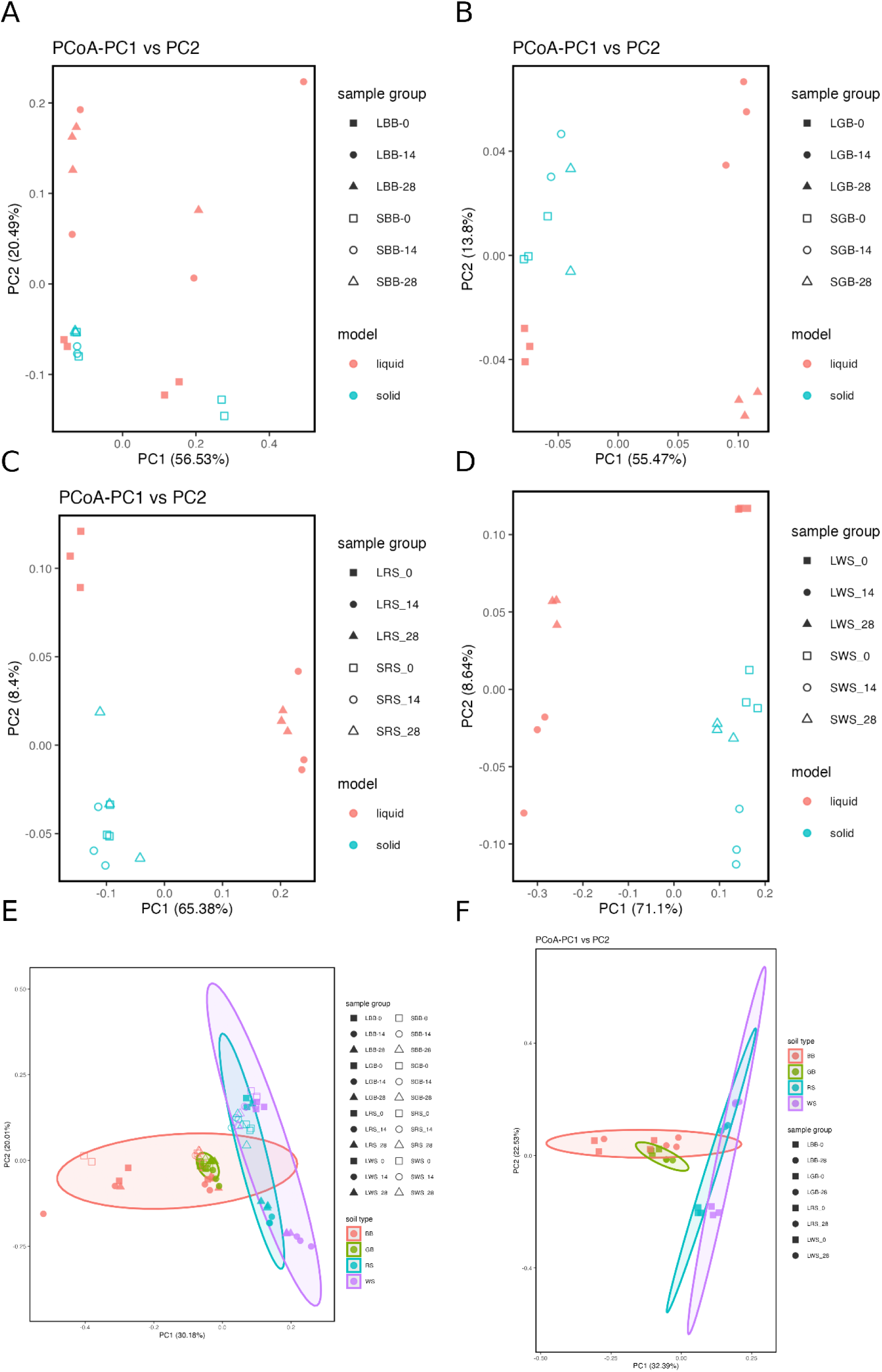
Beta diversity analysis of soil and NS-SESOM samples. (A-D) Principal component analysis (PCoA) of weighted UniFrac distances between sample conditions over time for black bag samples (BB, A), green bag samples (GB, B), roadside samples (RS, C) and wooded samples (WS, D). Blue open symbols represent solid soil samples and pink solid symbols represent liquid NS-SESOM at zero days (squares), 14 days (circles) and 28 days (triangles). (E-F) Comparative analysis by soil type weighted UniFrac distances. (E) PCoA of weighted UniFrac of all study samples. (F) PCoA of weighted UniFrac distances, comparing the day zero and day 28 NS-SESOM samples for each soil type.

To make comparisons between samples from different soil types, we next determined the Weighted UniFrac distances for all samples and all timepoints (Fig. 4E) and for the NS-SESOM samples at day zero and day 28 in culture for all soil types (Fig. 4F). In both instances, samples segregated along PC1 according to their soil type – commercial (pink and green symbols) or environmental (blue and purple symbols) – suggesting that the initial soil type remains the primary defining feature of NS-SESOM SMCs even after 28 days of liquid culture. The collective results of taxonomic, alpha diversity, and beta diversity analyses reveal that NS-SESOM preparations retain distinct, highly diverse, soil-specific SMC signatures over extended periods in liquid culture.

### Network analysis reveals conservation of community stability and complexity in NS-SESOM

The stability and complexity of an SMC is an emergent property of the relationships between individual members (Guseva et al., 2022; Wagg et al., 2019; Yuan et al., 2021). These relationships can be analyzed through microbial association (co-occurrence) network analyses. To compare the stability and complexity of NS-SESOM SMC networks to solid soils, we used NetCoMi (*<u>Net</u>*work *<u>Co</u>*nstruction and comparison for *<u>Mi</u>*crobiome data (Peschel et al., 2021)) to build microbial co-occurrence networks based on the top 50 most abundant ASVs by class (Fig. 5A) under each sample condition. Networks were constructed for the starting solid soil at Day 0 (Fig. 5B), and the NS-SESOM at Day 0 (Fig. 5C), Day 14 (Fig. 5D) and Day 28 (Fig. 5E). Visually, distinct similarities between the networks were observed. In general, positive associations were distributed within separate clusters, while negative relationships were observed to connect different clusters.

**Figure 5.**
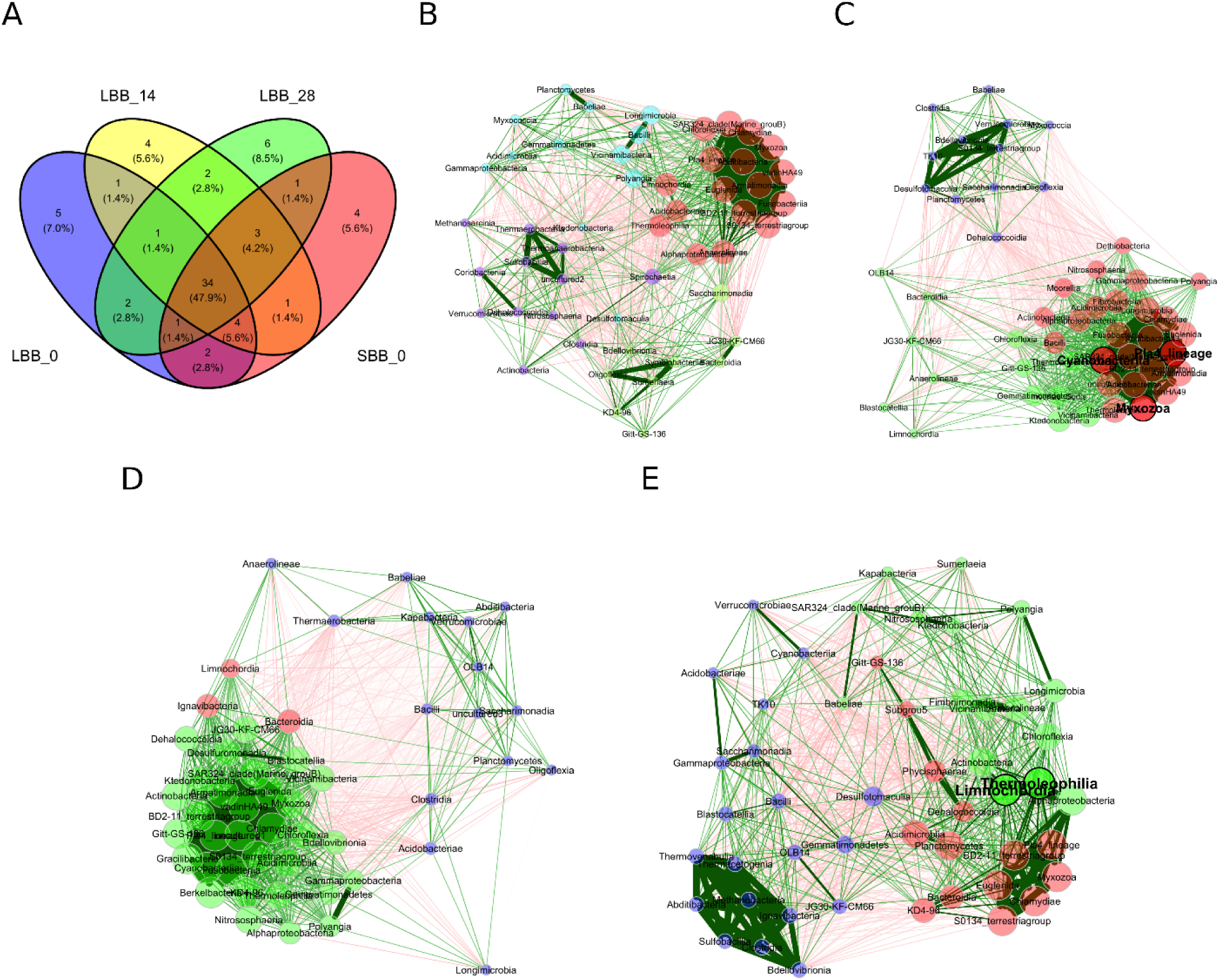
Network analysis of the top 50 most abundant ASVs for the BB commercial soil sample. A. Venn diagram showing the overlap of the top 50 most abundant ASVs at the class level for each condition from the starting solid soil (SBB_0), the initial liquid NS-SESOM extraction at day 0 (LBB_0), and the NS-SESOM after 14 days (LBB_14) and 28 days (LBB_28) in continuous culture. B. Class-level network from the starting solid soil (SBB_0). C. Class-level NS-SESOM network for the initial liquid at day 0 (LBB_0). (D) Class-level NS-SESOM network after 14 days. (LBB_14). (E) Class-level NS-SESOM network after 28 days. (LBB_28). Red edges represent negative associations while green edges represent positive associations. Node colors represent clusters. Node size is scaled by eigenvector centrality.

To quantify the comparison between networks, global network properties were calculated. Statistical analysis of global network properties including clustering coefficient, edge density, edge connectivity, and average dissimilarity revealed no statistically significant differences among any of the network properties examined for any of the BB Day 0, Day 14 or Day 28 NS-SESOM networks relative to the solid starting soil (Supplementary Table S11). The results strongly suggest a high level of conservation of major relationships between ASVs within the community over the 28-day liquid culture period.

The same network analysis and global network properties comparisons were performed for the three other soil types (GB, RS, and WS, Supplementary Figures S11-S13, Supplementary Table S11). Again, in all instances, there were no statistically significant differences in the network features of any of the Day 0, Day 14 or Day 28 NS-SESOM networks relative to the respective solid starting soil. The results overwhelmingly suggest that while there were significant changes in the relative abundance of some dominant taxa in different soils and NS-SESOM preparations based on LME analysis, the structural relationships of major taxa within the SMC networks are maintained in-tact over 28 days of continuous liquid culture.

### Soil microbial community functions are maintained over time in NS-SESOM

SMC composition dictates the higher-order functions of communities within soils (Hartmann & Six, 2023; Lee et al., 2025; Philippot et al., 2024b). To assess SMC function, we performed ASV functional pathway annotations with PICRUSt2 (Douglas et al., 2020) (Supplementary Fig. S14, Supplementary Table S12). The diversity of pathways represented in solid soil versus liquid NS-SESOM communities was generally highly conserved. Pathway alpha diversity was highly stable between solid soil and NS-SESOM samples in the rich commercial potting soils, whereas pathway alpha diversity increased over time in the liquid NS-SESOM samples relative to soil soils for the two environmental samples (Supplementary Figure S15), suggesting possible adaptation of communities to the resource-limited liquid environment. Pathway beta diversity was similarly conserved, and as noted for the taxonomic analysis, was primarily dictated by the starting soil type (Supplementary Figure S16).

Functional pathway networks for BB samples were built with NetCoMi from the 50 most abundant functional pathways under the varying different soil conditions. We observed that functional pathways were even more highly conserved between soil and NS-SESOM networks than ASVs, with 34 of 50 (68%) of the top 50 ASVs by class overlapping among the four networks (Fig. 5A) whereas 47 of 50 (94%) of the top 50 functional pathways overlapped among the four networks (Fig. 6A). Again, we observed high conservation of network architectures over the 28-day NS-SESOM culture period, relative to the starting soil (Fig. 6B-E). No statistically significant differences exist between any of the global network features for any of the pathway networks (Supplementary Table S13). These observations were fully recapitulated for the GB, RS and WS soil types (Supplementary Figs. S17 – S19). The collective results suggest that modest but significant changes in the NS-SESOM community composition ASVs over time in culture do not significantly disrupt the overall functions of the SMCs.

**Figure 6.**
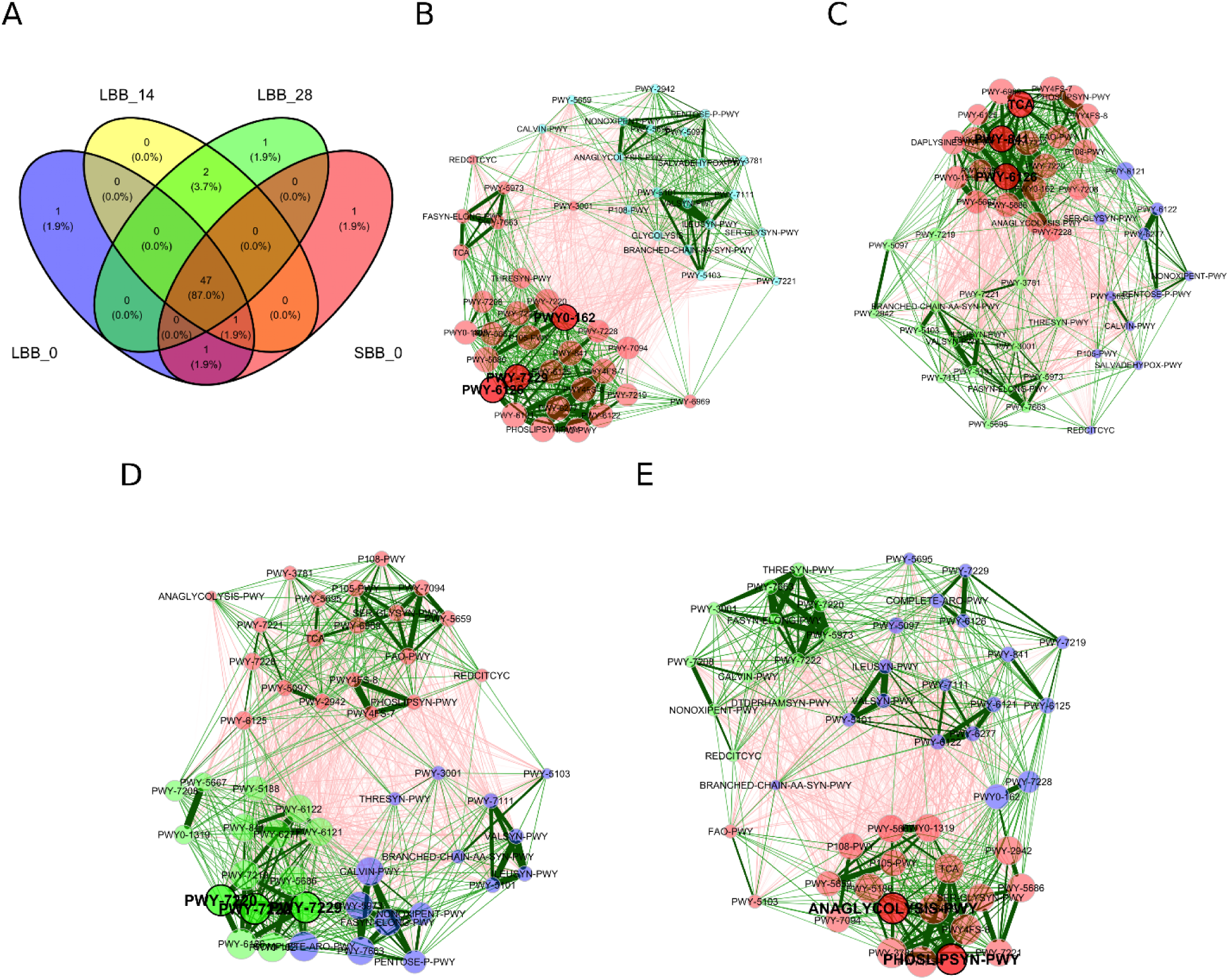
Network analysis of the top 50 most annotated functions for the BB commercial soil sample. A. Venn diagram showing the overlap of the top 50 most annotated functions for each condition from the starting solid soil (SBB_0), the initial liquid NS-SESOM extraction at day 0 (LBB_0), and the NS-SESOM after 14 days (LBB_14) and 28 days (LBB_28) in continuous culture. B. Pathway-level network from the starting solid soil (SBB_0). C. Pathway-level NS-SESOM network for the initial liquid at day 0 (LBB_0). (D) Pathway-level NS-SESOM network after 14 days. (LBB_14). (E) Pathway-level NS-SESOM network after 28 days. (LBB_28). Red edges represent negative associations while green edges represent positive associations. Node colors represent clusters. Node size is scaled by eigenvector centrality.

### The SESOM platform can be used to measure survival dynamics of an engineered microbe

We next tested whether our ST-SESOM and NS-SESOM platform could be used to measure the survival of an engineered organism within a complex native SMC. We used a 24-well deep well plate format to grow the soil microbe *P. putida* KT2440 expressing tdTomato (*P. putida* + tdTomato) as a model engineered organism in ST-SESOM and NS-SESOM preparations from the commercial BB soil, and measured *P. putida* survival by flow cytometry (Fig. 7, Supplementary Fig. S20-S21). In BB soil, *P. putida* persisted stably without measurable growth over 14 days in ST-SESOM but declined rapidly to undetectable levels by day 6 in NS-SESOM (Fig. 7A), suggesting that *P. putida* is not competitive with the existing SMC in this soil sample but can subsist in the nutritional environment of this carbon-rich soil. As a reference, we performed a traditional survival assay in BB soil using the colony forming unit assay (CFU, Fig. 7B). *P. putida* + tdTomato inoculated into autoclave sterilized soil grew exponentially and established stable colonization of the soil. Consistent with prior reports (Lázaro et al., 1998; Molina et al., 2000), *P. putida* + tdTomato inoculated into nonsterile soil showed no growth and a modest population decline over time, indicative of competition with the native SMC. Notably, by day 6, both assays were able to reliably differentiate between sterilized and nonsterile samples based on the relative engineered *P. putida* population. These results demonstrate SESOM models can be used in conjunction with high-throughput flow cytometry in a multi-well plate format to measure the performance of an engineered organism within a native SMC in a way that recapitulates relevant survival dynamics.

**Figure 7.**
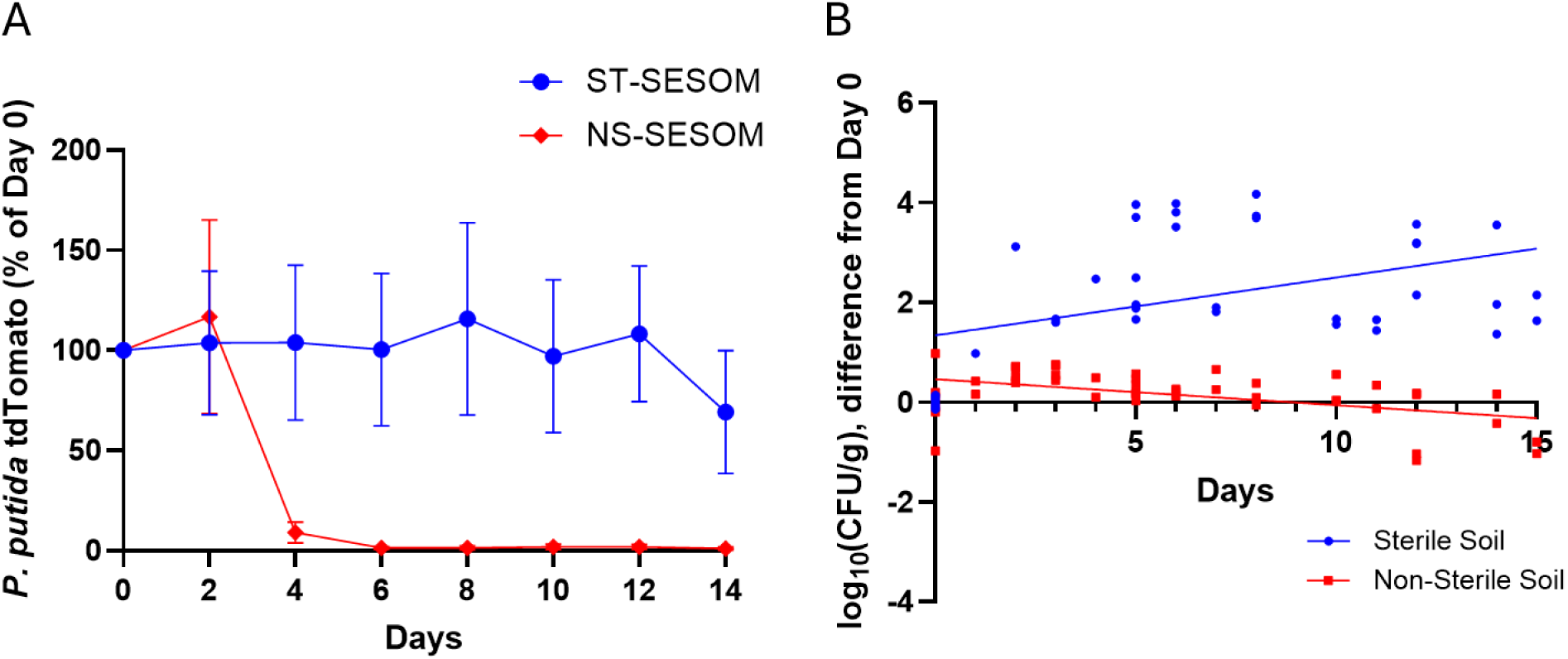
Survival of *P. putida* expressing tdTomato in SESOM and solid soil. (A) Sterile (ST) and non-sterile (NS) SESOM preparations were made with BB commercial soil. *P. putida* expressing tdTomato was inoculated into SESOM and sampled by flow cytometry at timepoints indicated. Error bars are SEM (n=3). (B) *P. putida* expressing tdTomato was inoculated into 10g of autoclave sterilized (sterile) or non-sterile BB commercial soil in a glass petri dish, and sampled by colony forming unit assay (CFU) at timepoints indicated. Linear regression analysis is shown (n=7).

## DISCUSSION

The work presented here began with a simple question: how stable are liquid-extracted SMCs? Several prior studies with defined SynComs demonstrated community collapse, dominated by one or a few key taxa, over an extended time in liquid culture (Cira et al., 2018; Gilmore et al., 2019; Niu et al., 2017; Velte et al., 2025). However, some of these same studies noted comparatively stable behavior of native consortia (Gilmore et al., 2019; Velte et al., 2025). Our results corroborate the significant stability of native, NS-SESOM-extracted SMCs over a 28-day period in liquid culture (Fig. 2, Fig. 5). While SMCs do undergo evolution over time in liquid culture with a minor loss in alpha diversity (Fig. 3), they retain key microbial signatures associated with their starting soil types (Fig. 4). More importantly, network analysis of functional pathways associated with the dominant taxa shows that the function of the NS-SESOM SMCs is entirely conserved throughout a 28-day culture protocol (Fig. 6).

We observed that in all NS-SESOM sample types, the Day 0 – Day 14 time period shows greater shifts in the community structure than the Day 14 – Day 28 time period; the solid Day 0 and NS-SESOM Day 0 were highly similar in all cases. There are two implications of these observations: First, that short term experiments are likely to faithfully reflect the solid community dynamics for a brief period of time, though the exact window will need to be experimentally validated; Second, at some point between within the first 14 days of culture, the community stabilized into a liquid-adapted structure that retained the key taxa signature (Fig. 4) and importantly, the community function (Fig. 6) of the initial starting soil. It may be desirable to permit this stabilization in some experimental contexts. Additional experiments to map the evolution of the community over time and in a specific soil context would be needed.

We noted a correlation between the stability of the taxa (Fig. 2) and the organic content of the soil (Supplementary Table 1). The high organic matter content of the NS-SESOM communities from the commercial potting soils (∼50-70%) were more stable over time than the environmental samples (∼2-15%). Although it is noted (as discussed above) that all samples, regardless of the soil physicochemical properties, settled into stable community structures between days 14 and 28 of culture. An obvious hypothesis would be that the availability of carbon sources decreases competition between species and stabilizes the community structure. However, some reports suggest that nutrient limitation may in some contexts be a stabilizing factor for microbial communities (Ratzke et al., 2020; Ye et al., 2026). Our observations raise the possibility that carbon source supplementation may be manipulated to modulate the community stability, though this hypothesis remains to be tested. We specifically chose PBS for extraction, to help stabilize pH while eliminating a potential carbon source (MOPS) that could be metabolized by *P. putida* or other community members (Supplementary Fig. S1) because we wanted to understand the dynamics within the existing resource-limited context of the soil. Future experiments could investigate the role of carbon availability in community stability using our NS-SESOM system.

Finally, we demonstrate that the survival and persistence dynamics of an engineered microbe, the synthetic biology environmental chassis *P. putida*, in a soil environment, can be recapitulated in NS-SESOM (Fig. 7). There is a need for a variety of synthetic soil models with varying levels of complexity to facilitate the development of engineered microbial solutions (Orebaugh et al., 2026; Zhalnina et al., 2018). Our NS-SESOM model provides an additional platform between laboratory culture conditions and field soil samples that captures many of the microbial and chemical properties of a specific target soil, while maintaining the flexibility and high-throughput facility of a liquid culture.

## CONCLUSION

The NS-SESOM model is a validated, reproducible liquid SMC model. The taxonomic composition and diversity analyses reveal NS-SESOM SMCs are highly reproducible, faithful to the initial SMC, and stable over extended time in culture up to 28 days. Despite some shifts in SMC composition over extended time in culture, the NS-SESOM SMCs maintain unique signatures of their initial SMCs and their dominant functions. The platform also maintains members that are unculturable in isolation, and is amenable to high-throughput multi-well plate formats. Thus, we conclude that the NS-SESOM method of culturing liquid SMCs as a model of the soil microbiome is a useful platform with specific benefits over the bespoke assembly of liquid SynComs from cultured members. The implementation of the synthetic biology DBTL (design-build-test-learn) cycle requires flexible, modular, high-throughput assay platforms. We envision that NS-SESOM can be used in the future for the high-throughput assessment of engineered bacteria for a variety of soil applications including nitrogen fixation, bioremediation, and biomining.

## Data Availability

Raw data files in FASTQ format were deposited in the NCBI sequence read archive under accession number PRJNA1309165. All other raw data is available upon request.

## Acknowledgements

We thank Dr. Adam Guss, Dr. Joshua Elmore, and Dr. Jay Huenemann for the kind gift of the *P. putida* strain AG4775 and plasmids for the Bxb recombinase system, and for their protocols and helpful advice. This publication was developed in part under Assistance Agreement No. RD-84020601 awarded by the U.S. Environmental Protection Agency to N.G.F. It has not been formally reviewed by EPA. The views expressed in this document are solely those of the authors and do not necessarily reflect those of the Agency. EPA does not endorse any products or commercial services mentioned in this publication. We gratefully acknowledge funding support from the DARPA CERES (D24AC00011) and DARPA BioReporters (HR001119C0107) programs to N.G.F, and new faculty start-up funds from Worcester Polytechnic Institute to N.G.F.

## Disclosures

– The authors declare no competing interests.

## Author Contributions

S.L. performed all bioinformatics analyses and related data curation and visualizations. G.N.C.P and S.V. optimized the SESOM extraction methodology. G.N.C.P. performed DNA extractions for 16S sequencing, and flow cytometry analyses and related visualizations. Bacterial culture and CFU experiments were performed by N.G.F., S.V., N.K., and A.S. S.L. wrote the initial draft of the manuscript. N.G.F edited the manuscript. All authors reviewed and approved of the final draft. N.G.F. conceived of the project, acquired funding support, provided project management and supervision.

## Notes

### Competing Interest Statement

The authors have declared no competing interest.

