## Supplementary Figures for "Long-term stability of soil microbiome structure and function in liquid soil extracts"

The Supplementary Information accompanying this manuscript contains 21 Supplementary Figures (S1-S21), and 13 Supplementary Tables (S1-S13).

Supplementary Figure S1

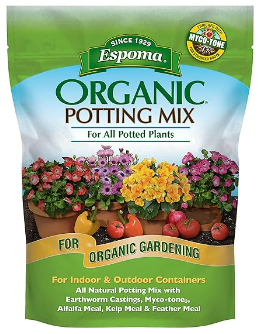

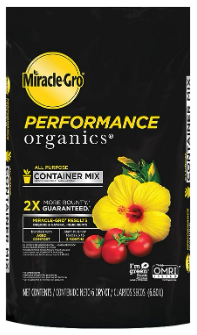

A. Black Bag Sample

B. Green Bag Sample

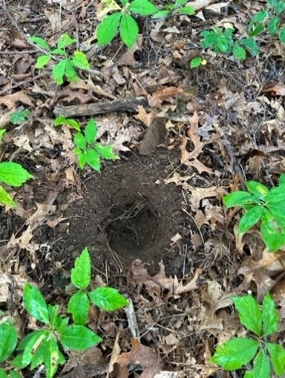

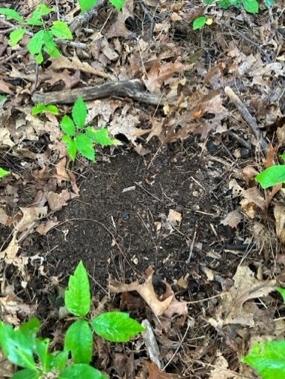

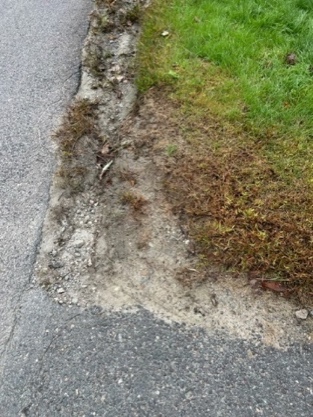

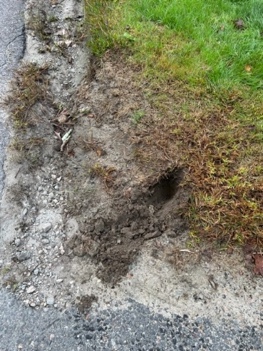

**DNA extraction**

D. Roadside Sample

C. Wooded Sample

**Supplementary Figure S1. Soil samples.** Commercial samples are (A) Miracle-Gro Performance Organics Container Mix (Black bag sample, BB) and (B) Espoma Organic Potting Mix (Green bag sample, GB). Environmental samples are (C) Wooded sample (WS) and (D) Roadside sample (RS).

Supplementary Figure S2

B.

A.

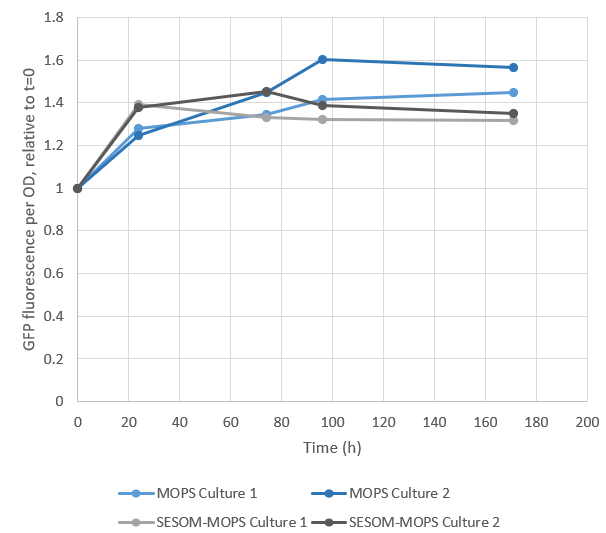

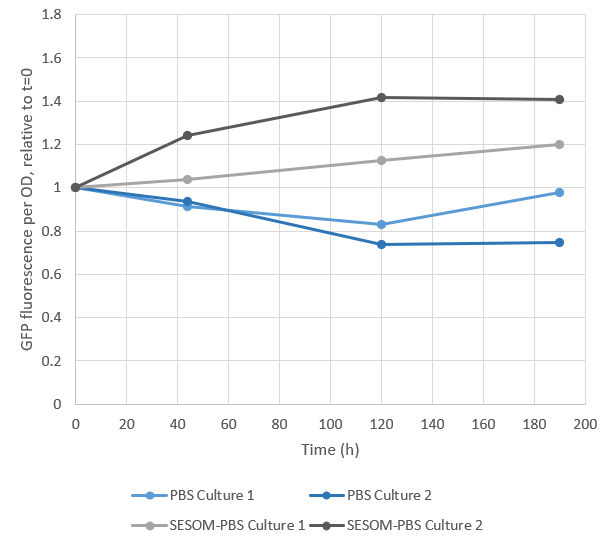

**Supplementary Figure S2. Comparison of SESOM extraction in MOPS and PBS**. *P. putida* expressing mNeonGreen was cultured in MOPS buffer or SESOM made with MOPS (A), or PBS or SESOM made with PBS (B). Fluorescence (excitation 485 nm, emission 535 nm) and optical density (OD) at 600 nm measurements were made on a Perkin Elmer NIVO plate reader at the times indicated. Fluorescence per OD measurements are shown relative to the time zero condition. Two independent replicate cultures are shown for each condition. These preliminary results indicate that *P. putida*, which is capable of complex catabolism, may possibly catabolize MOPS.

Supplementary Figure S3

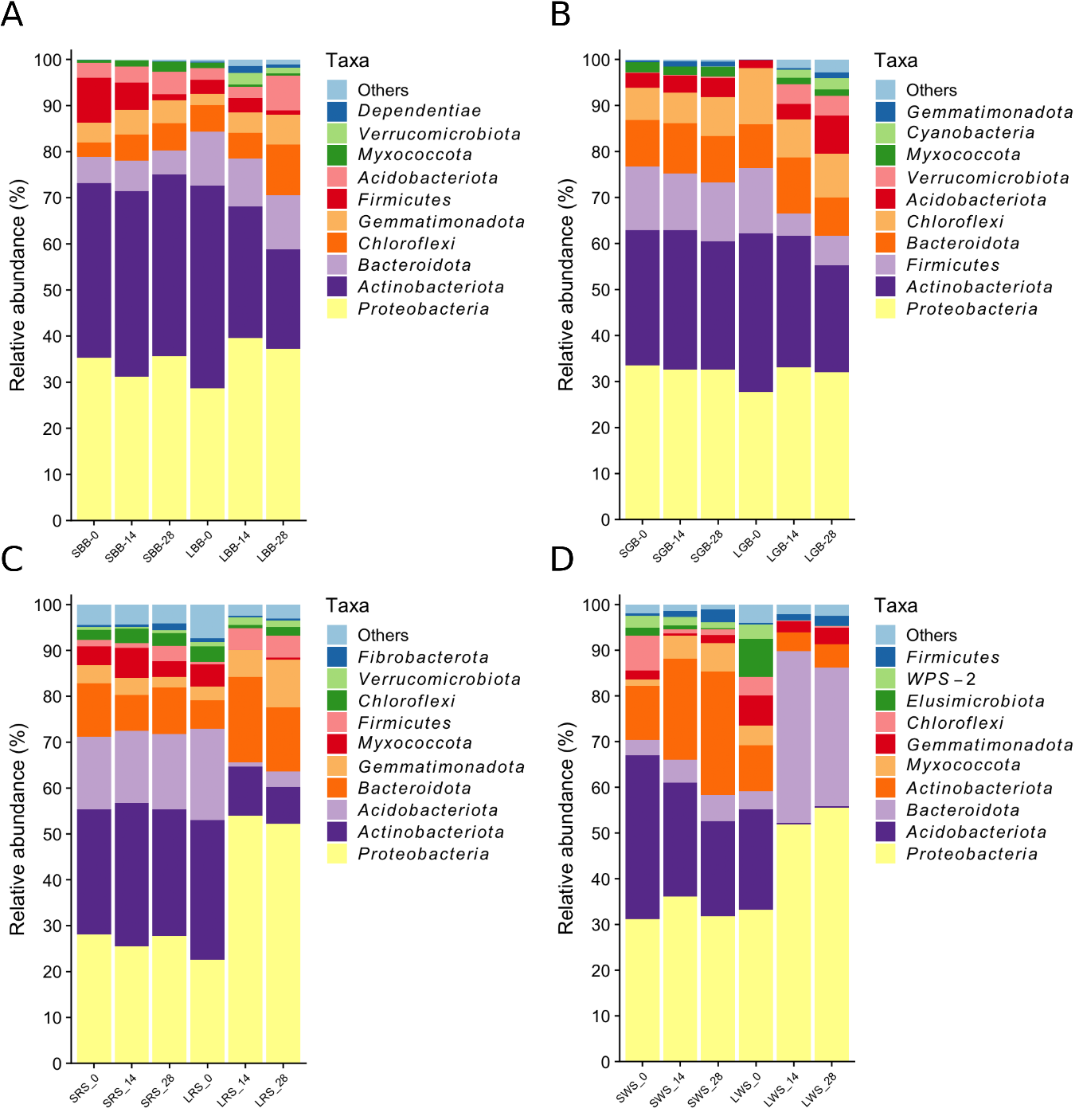

**Supplementary Figure S3. Relative abundance of bacterial taxa by phylum in solid soils and NS-SESOM**. Sample ID tags are assembled as follows: L (liquid) represents NS-SESOM and S represents solid soil samples; commercial samples are BB (black bag) and GB (green bag), environmental samples are RS (roadside) and WS (wooded). The DNA samples were collected at day 0, day 14 and day 28, as indicated by the number following each sample ID tag, and subjected to 16S sequencing and ASV annotations by phylum. The 10 most frequent bacteria phyla are indicated by colors.

Supplementary Figure S4

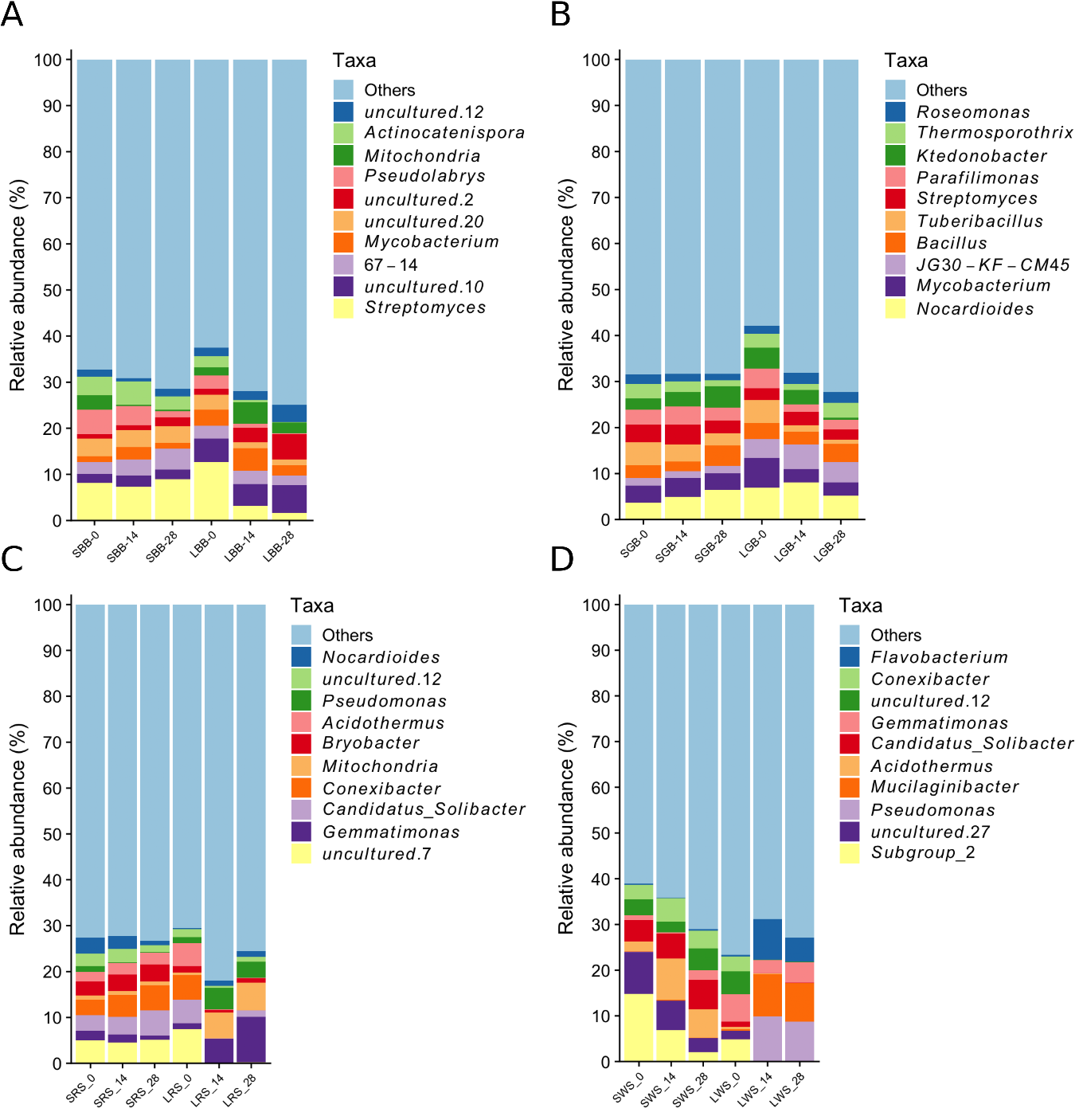

**Supplementary Figure S4. Relative abundance of bacterial taxa by genus in solid soils and NS-SESOM**. Sample ID tags are assembled as follows: L (liquid) represents NS-SESOM and S represents solid soil samples; commercial samples are BB (black bag) and GB (green bag), environmental samples are RS (roadside) and WS (wooded). The DNA samples were collected at day 0, day 14 and day 28, as indicated by the number following each sample ID tag, and subjected to 16S sequencing and ASV annotations by genus. The 10 most frequent bacteria genera are indicated by colors.

Supplementary Figure S5

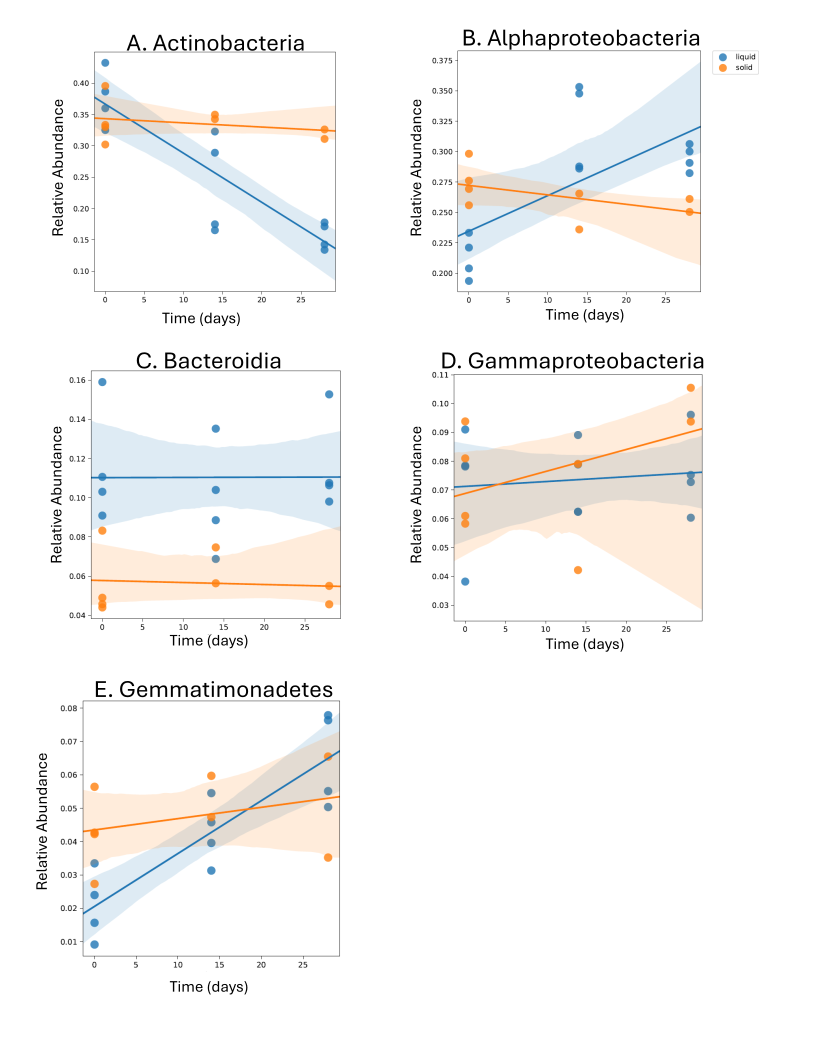

**Supplementary Figure S5. Linear mixed effects (LME) models of the top five most abundant taxa by class in BB soil**. LME models are shown for the top five classes at sample days zero, 14 and 28 in order of abundance: Actinobacteria (A), Alphaproteobacteria (B), Bacteroidia (C), Gammaproteobacteria (D), and Gemmatimonadetes (E). Solid soil sample replicates are shown in orange; NS-SESOM sample replicates are shown in blue.

Supplementary Figure S6

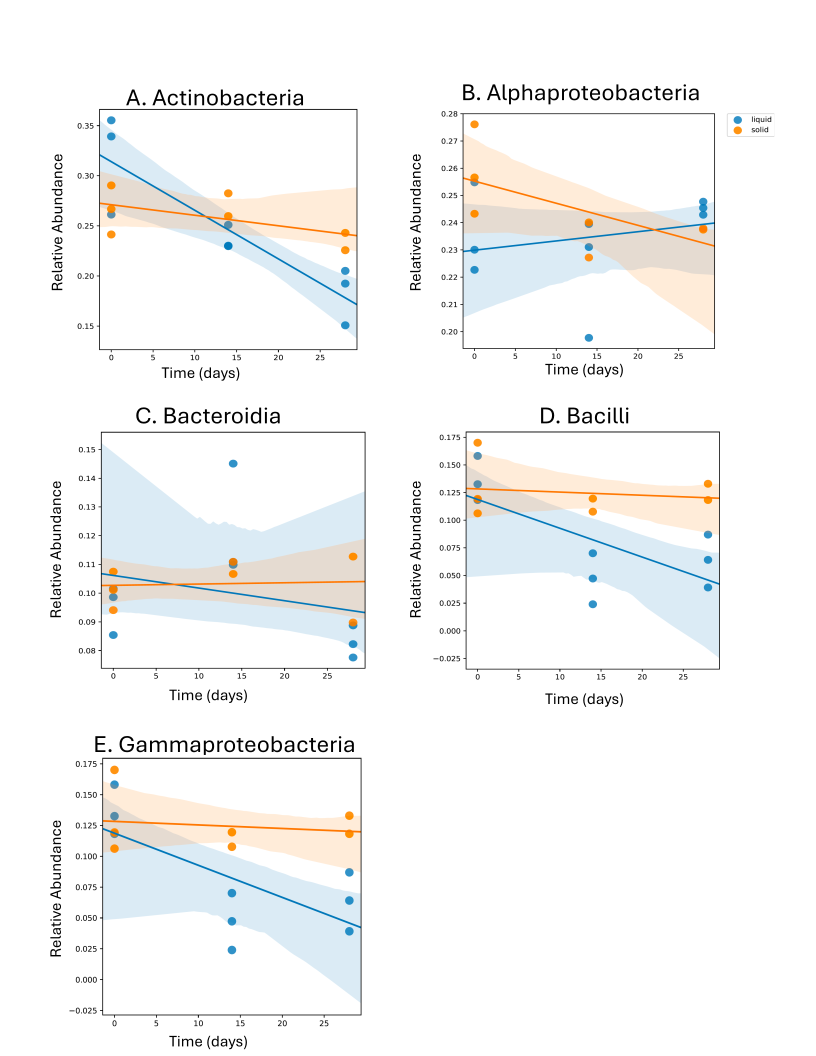

**Supplementary Figure S6. Linear mixed effects (LME) models of the top five most abundant taxa by class in GB soil**. LME models are shown for the top five classes at sample days zero, 14 and 28 in order of abundance: Actinobacteria (A), Alphaproteobacteria (B), Bacteroidia (C), Bacilli (D), and Gammaproteobacteria (E). Solid soil sample replicates are shown in orange; NS-SESOM sample replicates are shown in blue.

Supplementary Figure S7

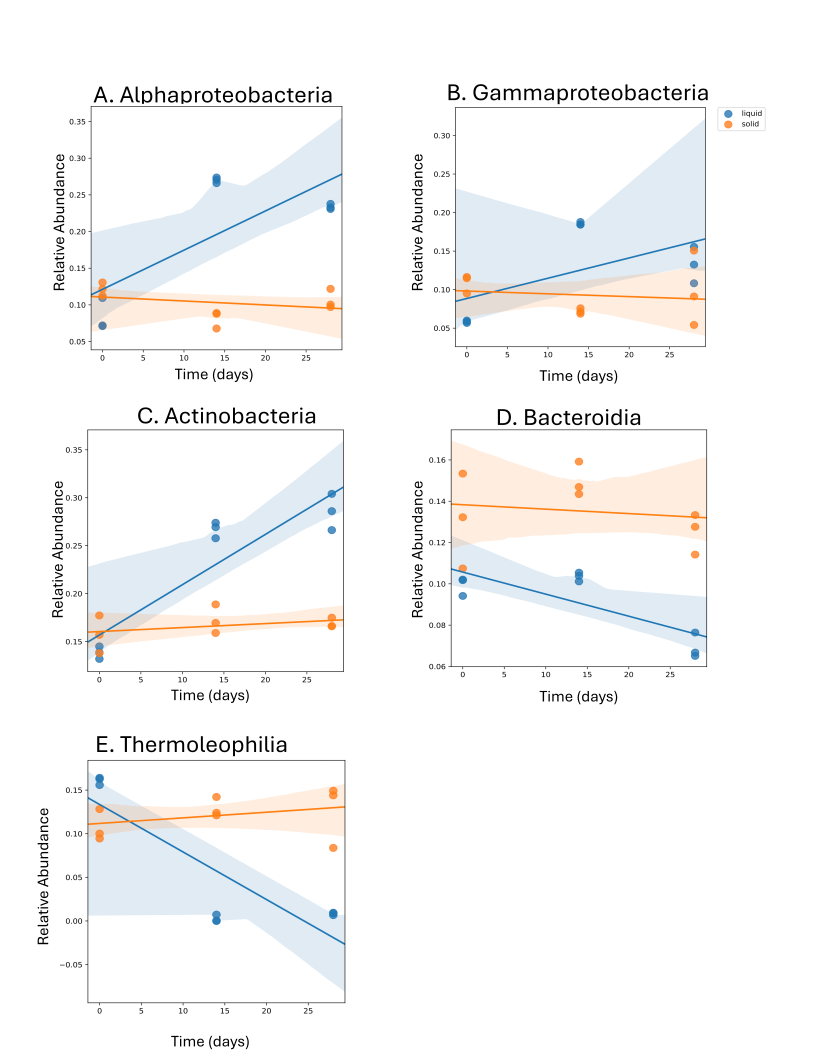

**Supplementary Figure S7. Linear mixed effects (LME) models of the top five most abundant taxa by class in RS soil**. LME models are shown for the top five classes at sample days zero, 14 and 28 in order of abundance: Alphaproteobacteria (A), Gammaproteobacteria (B), Actinobacteria (C), Bacteroidia (D), and Thermoleophilia (E). Solid soil sample replicates are shown in orange; NS-SESOM sample replicates are shown in blue.

Supplementary Figure S8

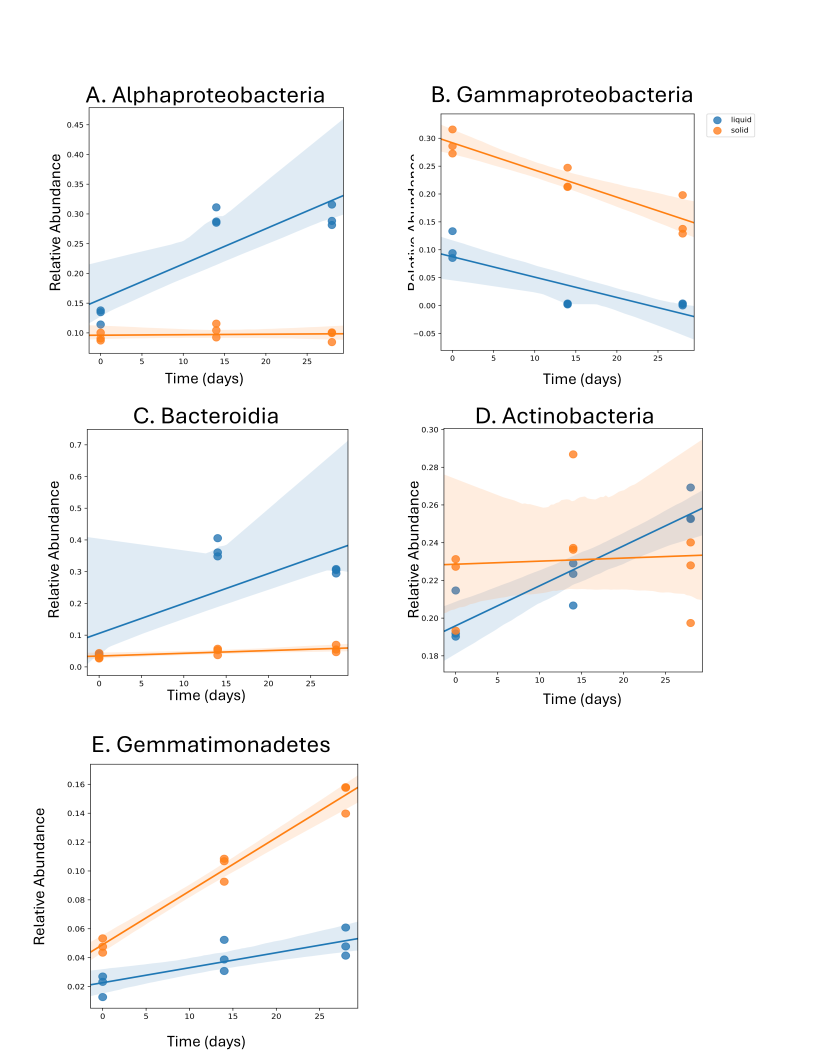

**Supplementary Figure S8. Linear mixed effects (LME) models of the top five most abundant taxa by class in WS soil**. LME models are shown for the top five classes at sample days zero, 14 and 28 in order of abundance: Alphaproteobacteria (A), Gammaproteobacteria (B), Bacteroidia (C), Actinobacteria (D), and Gemmatimonadetes (E). Solid soil sample replicates are shown in orange; NS-SESOM sample replicates are shown in blue.

Supplementary Figure S9

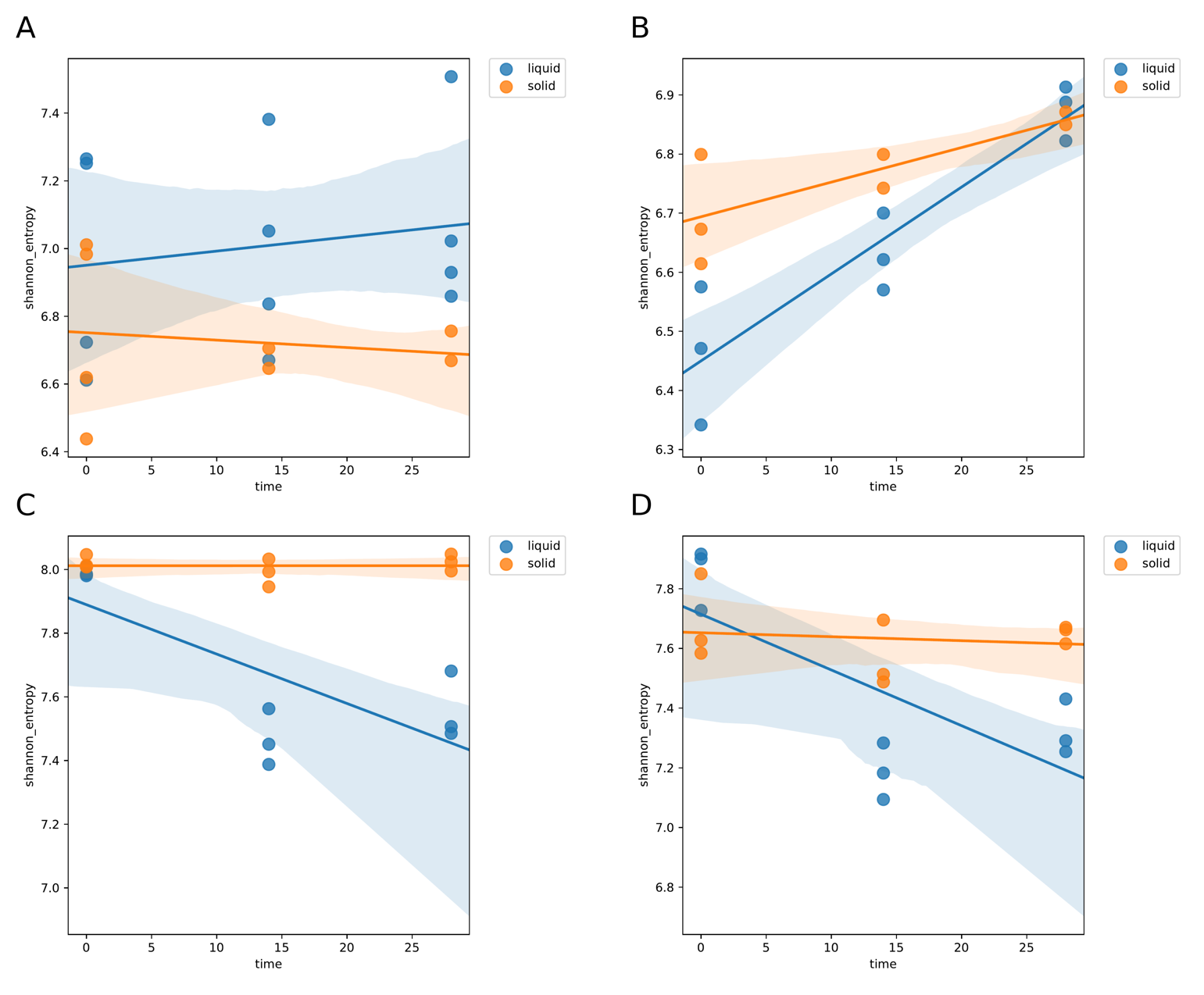
**Supplementary Figure S9. Linear mixed effects models for sample alpha diversity**. A. BB soil and NS-SESOM. B. GB soil and NS-SESOM. C. RS soil and NS-SESOM. D. WS soil and NS-SESOM. Solid soil sample replicates are shown in orange; NS-SESOM sample replicates are shown in blue.

Supplementary Figure S10

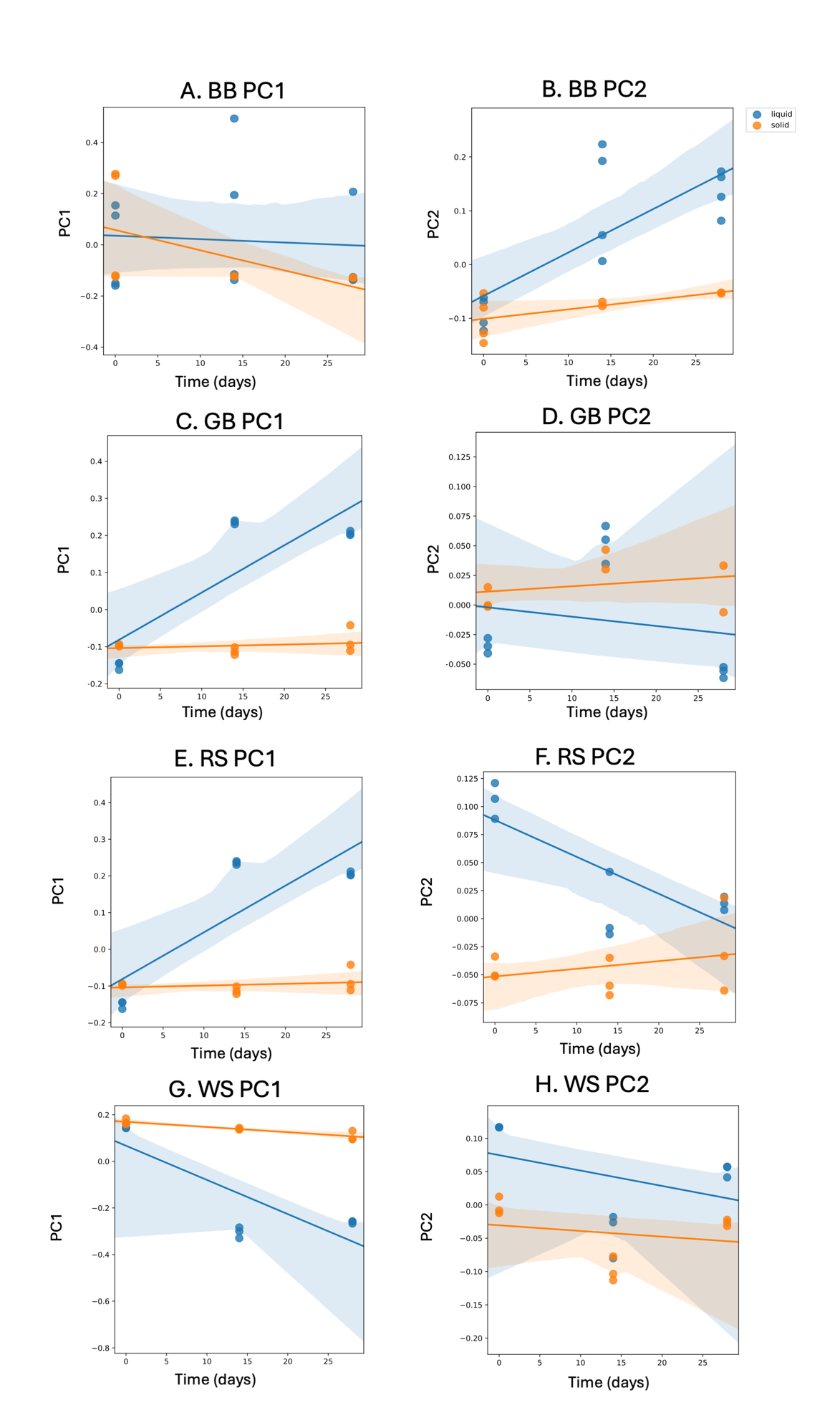

**Supplementary Figure S10. Linear mixed effects models for the first two principal components (PC1 and PC2) from Weighted UniFrac distances**. A-B. PC1 and PC2 from BB soil and NS-SESOM. C-D. PC1 and PC2 from GB soil and NS-SESOM. E-F. PC1 and PC2 from RS soil and NS-SESOM. G-H. PC1 and PC2 from WS soil and NS-SESOM. Solid soil sample replicates are shown in orange; NS-SESOM sample replicates are shown in blue.

Supplementary Figure S11

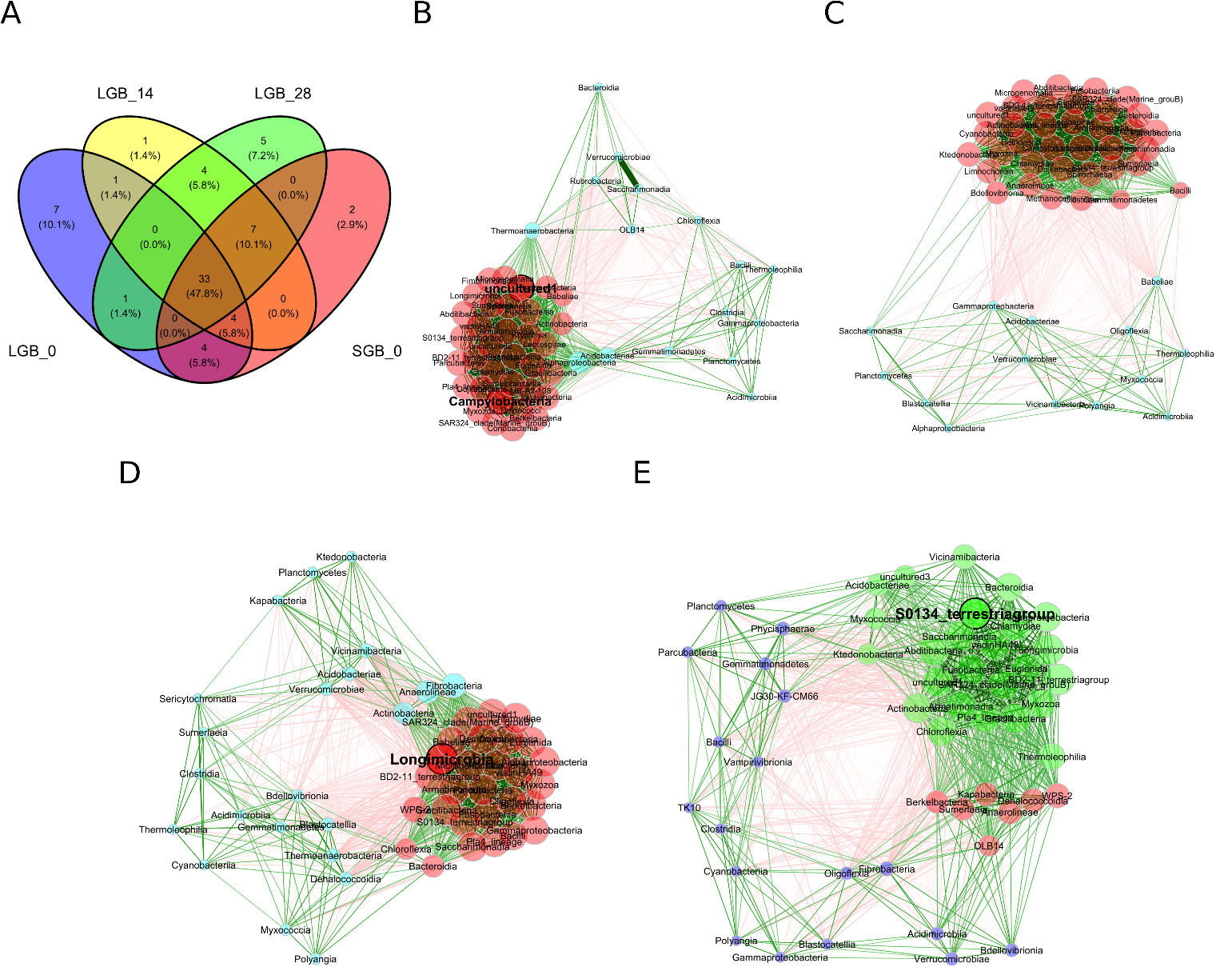

**Supplementary Figure S11. Network analysis of the top 50 most abundant ASVs for the GB commercial soil sample**. A. Venn diagram showing the overlap of the top 50 most abundant ASVs at the class level for each condition from the starting solid soil (SGB_0), the initial liquid NS-SESOM extraction at day 0 (LGB_0), and the NS-SESOM after 14 days (LGB_14) and 28 days (LGB_28) in continuous culture. B. Class-level network from the starting solid soil (SGB_0). C. Class-level NS-SESOM network for the initial liquid at day 0 (LGB_0). (D) Class-level NS-SESOM network after 14 days. (LGB_14). (E) Class-level NS-SESOM network after 28 days. (LGB_28). Red edges represent negative associations while green edges represent positive associations. Node colors represent clusters. Node size is scaled by eigenvector centrality.

Supplementary Figure S12

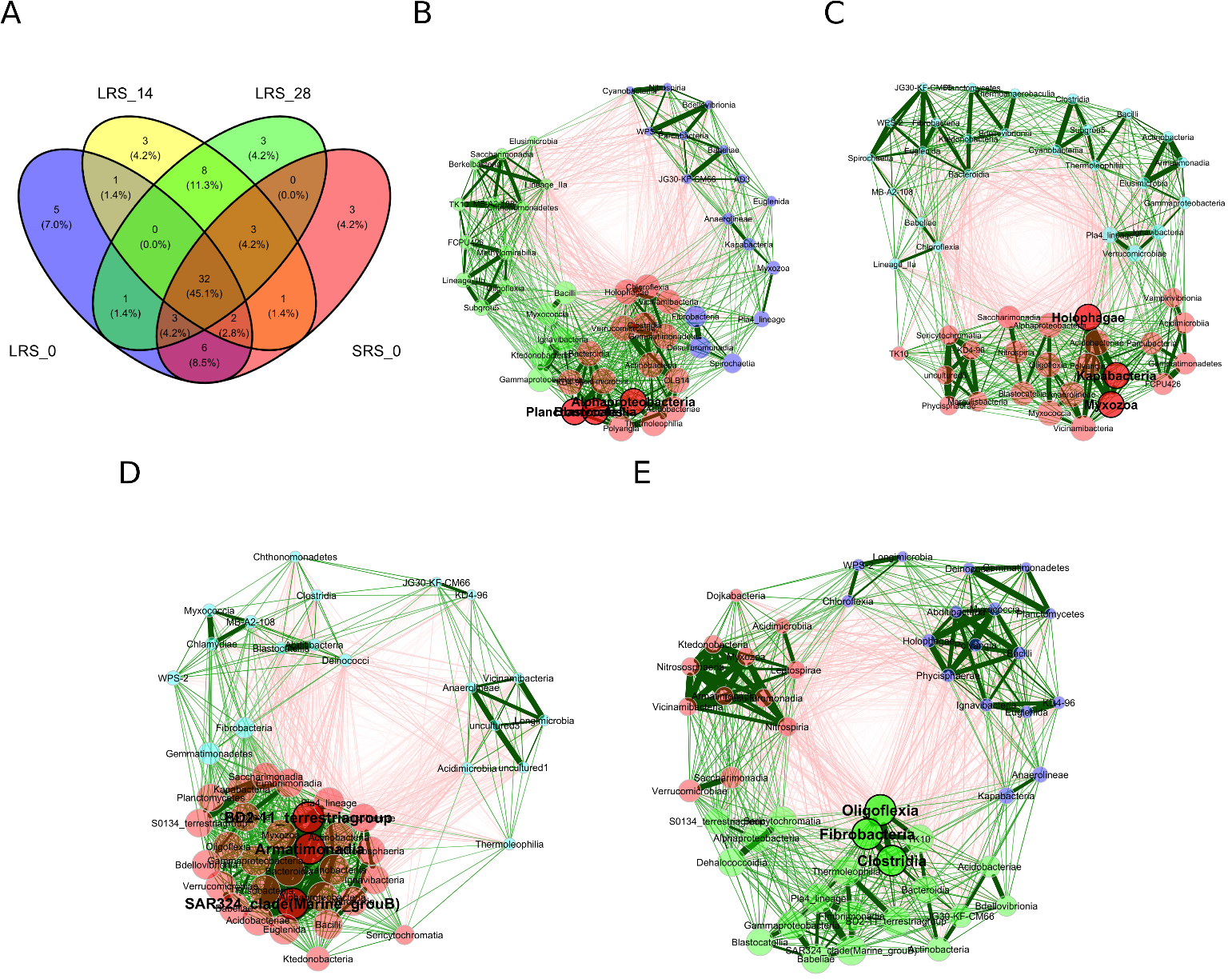

**Supplementary Figure S12.** **Network analysis of the top 50 most abundant ASVs for the RS environmental soil sample**. A. Venn diagram showing the overlap of the top 50 most abundant ASVs at the class level for each condition from the starting solid soil (SRS_0), the initial liquid NS-SESOM extraction at day 0 (LRS_0), and the NS-SESOM after 14 days (LRS_14) and 28 days (LRS_28) in continuous culture. B. Class-level network from the starting solid soil (SRS_0). C. Class-level NS-SESOM network for the initial liquid at day 0 (LRS_0). (D) Class-level NS-SESOM network after 14 days. (LRS_14). (E) Class-level NS-SESOM network after 28 days. (LRS_28). Red edges represent negative associations while green edges represent positive associations. Node colors represent clusters. Node size is scaled by eigenvector centrality.

Supplementary Figure S13

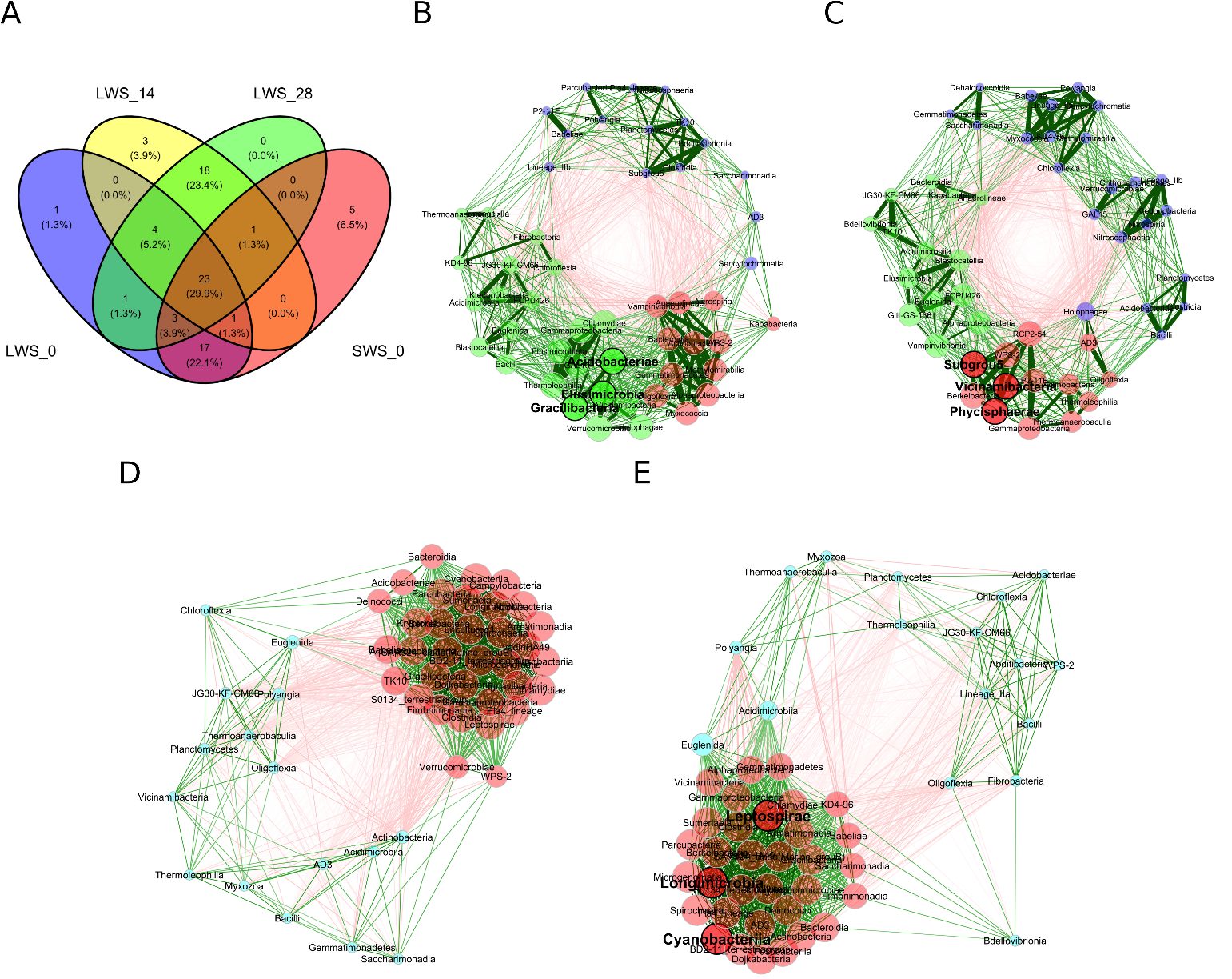

**Supplementary Figure S13. Network analysis of the top 50 most abundant ASVs for the WS environmental soil sample**. A. Venn diagram showing the overlap of the top 50 most abundant ASVs at the class level for each condition from the starting solid soil (SWS_0), the initial liquid NS-SESOM extraction at day 0 (LWS_0), and the NS-SESOM after 14 days (LWS_14) and 28 days (LWS_28) in continuous culture. B. Class-level network from the starting solid soil (SWS_0). C. Class-level NS-SESOM network for the initial liquid at day 0 (LWS_0). (D) Class-level NS-SESOM network after 14 days. (LWS_14). (E) Class-level NS-SESOM network after 28 days. (LWS_28). Red edges represent negative associations while green edges represent positive associations. Node colors represent clusters. Node size is scaled by eigenvector centrality.

Supplementary Figure S14

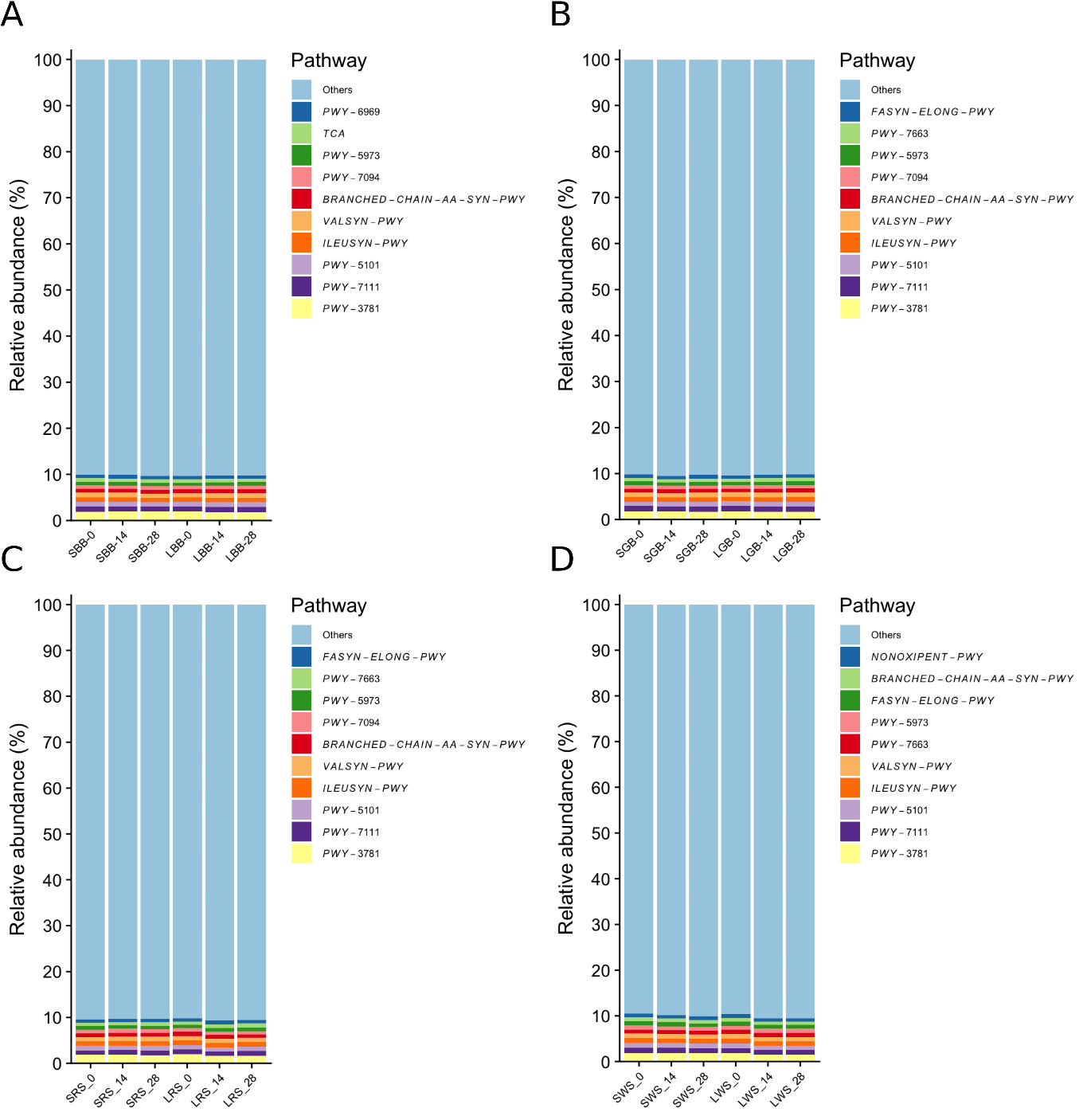

**Supplementary Figure S14. Top 10 pathway annotations in each sample.** A. BB soil (SBB) and NS-SESOM (LBB). B. GB soil (SGB) and NS-SESOM (LGB). C. RS soil (SRS) and NS-SESOM (LRS). D. WS soil (SWS) and NS-SESOM (LWS). Numbers for each sample ID tag indicate culture time in days.

Supplementary Figure S15

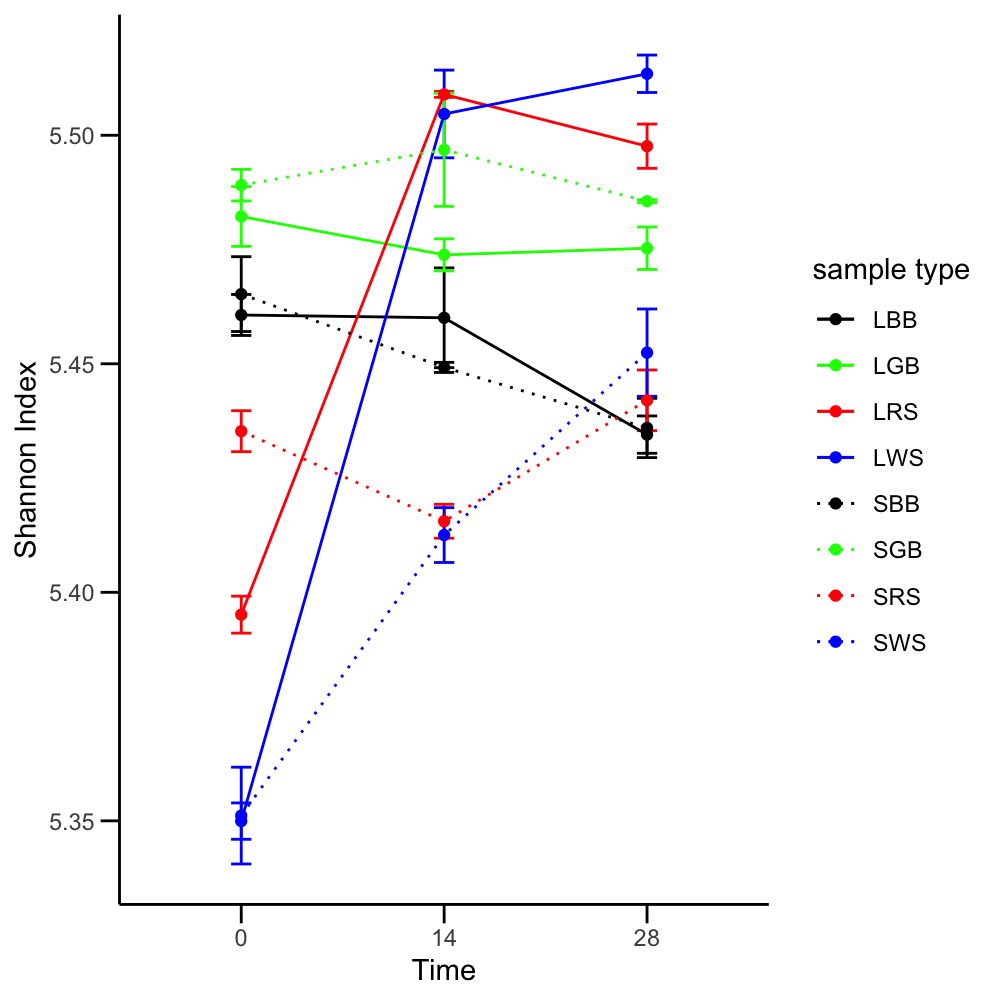

**Supplementary Figure S15.** **Alpha diversity indices of pathway annotations from soils and NS-SESOM samples.** Sample ID tags are assembled as follows: L (liquid) represents NS-SESOM (solid lines), and S represents solid soil samples (dashed lines); commercial samples are BB (black bag) and GB (green bag), environmental samples are RS (roadside) and WS (wooded sample). ASV annotations from 16S sequence analysis were used to identify community functions using PICRUSt2. Shannon index was then calculated for each sample. Error bars represent SEM (n=3).

Supplementary Figure S16

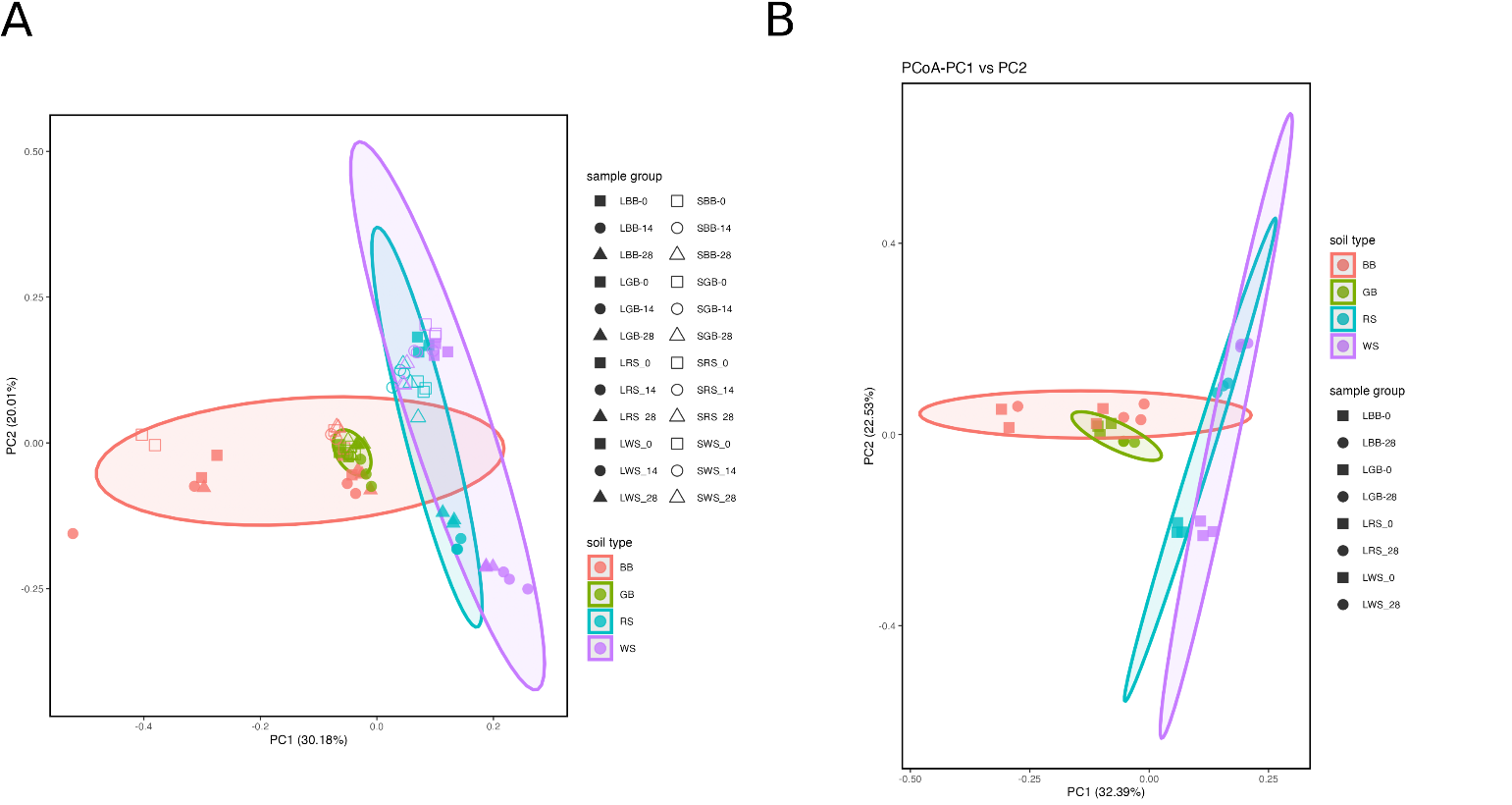

**Supplementary Figure S16. Principal component analysis (PCoA) of Bray-Curtis dissimilarity indices of pathway annotations between soil samples**. PCoA of Bray-Curtis dissimilarity analysis for all samples (A) and Day 0 versus Day 28 NS-SESOM samples only (B). Soil samples are indicated by color: BB (pink), GB (green), RS (blue) and WS (purple). Open symbols represent solid soil samples and solid symbols represent liquid NS-SESOM at zero days (squares), 14 days (circles) and 28 days (triangles).

Supplementary Figure S17

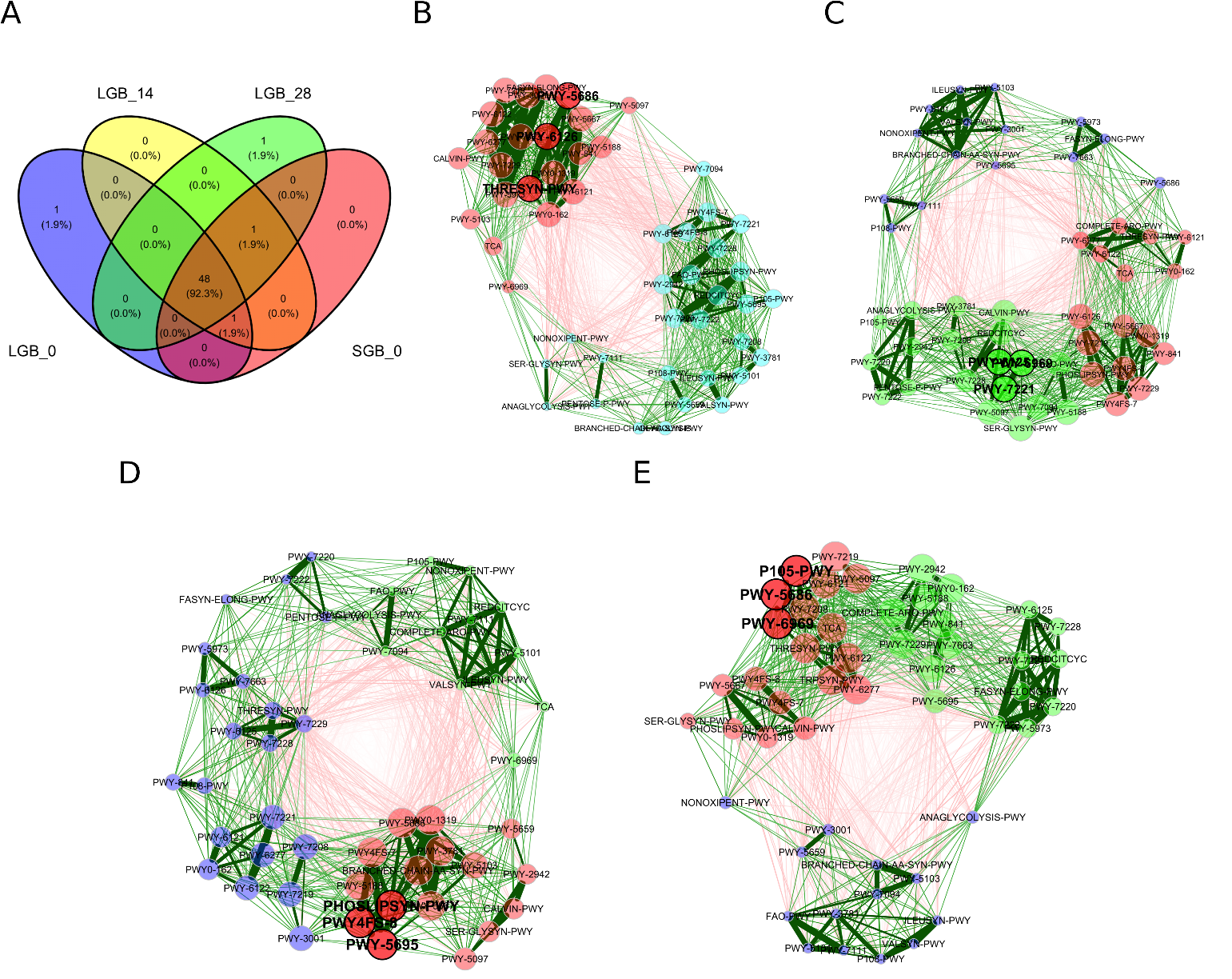

**Supplementary Figure S17. Network analysis of the top 50 most annotated functions for the GB commercial soil sample**. A. Venn diagram showing the overlap of the top 50 most annotated functions for each condition from the starting solid soil (SGB_0), the initial liquid NS-SESOM extraction at day 0 (LGB_0), and the NS-SESOM after 14 days (LGB_14) and 28 days (LGB_28) in continuous culture. B. Pathway-level network from the starting solid soil (SGB_0). C. Pathway-level NS-SESOM network for the initial liquid at day 0 (LGB_0). (D) Pathway-level NS-SESOM network after 14 days. (LGB_14). (E) Pathway-level NS-SESOM network after 28 days. (LGB_28). Red edges represent negative associations while green edges represent positive associations. Node colors represent clusters. Node size is scaled by eigenvector centrality.

Supplementary Figure S18

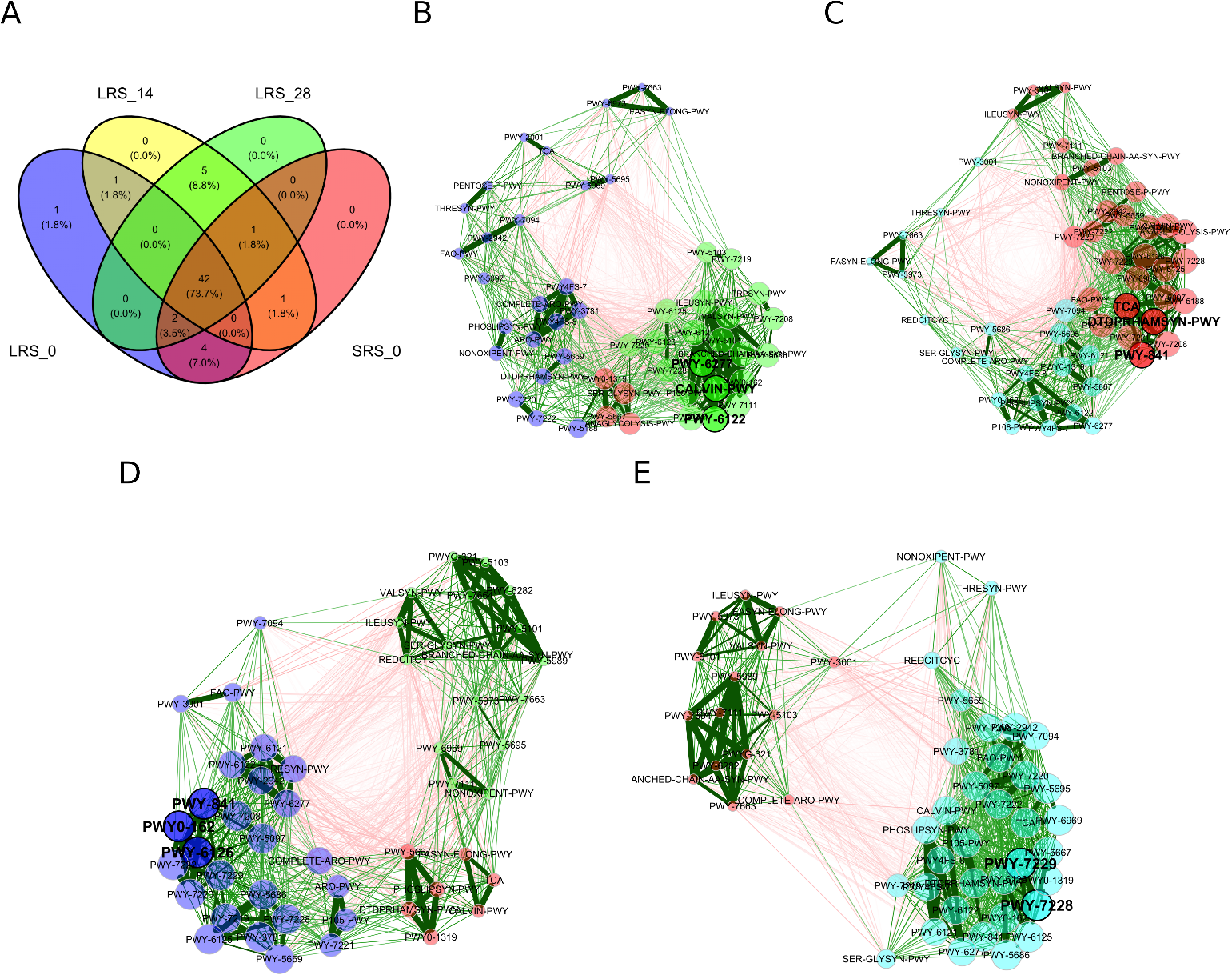

**Supplementary Figure S18.** **Network analysis of the top 50 most annotated functions for the RS environmental soil sample**. A. Venn diagram showing the overlap of the top 50 most annotated functions for each condition from the starting solid soil (SRS_0), the initial liquid NS-SESOM extraction at day 0 (LRS_0), and the NS-SESOM after 14 days (LRS_14) and 28 days (LRS_28) in continuous culture. B. Pathway-level network from the starting solid soil (SRS_0). C. Pathway-level NS-SESOM network for the initial liquid at day 0 (LRS_0). (D) Pathway-level NS-SESOM network after 14 days. (LRS_14). (E) Pathway-level NS-SESOM network after 28 days. (LRS_28). Red edges represent negative associations while green edges represent positive associations. Node colors represent clusters. Node size is scaled by eigenvector centrality.

Supplementary Figure S19

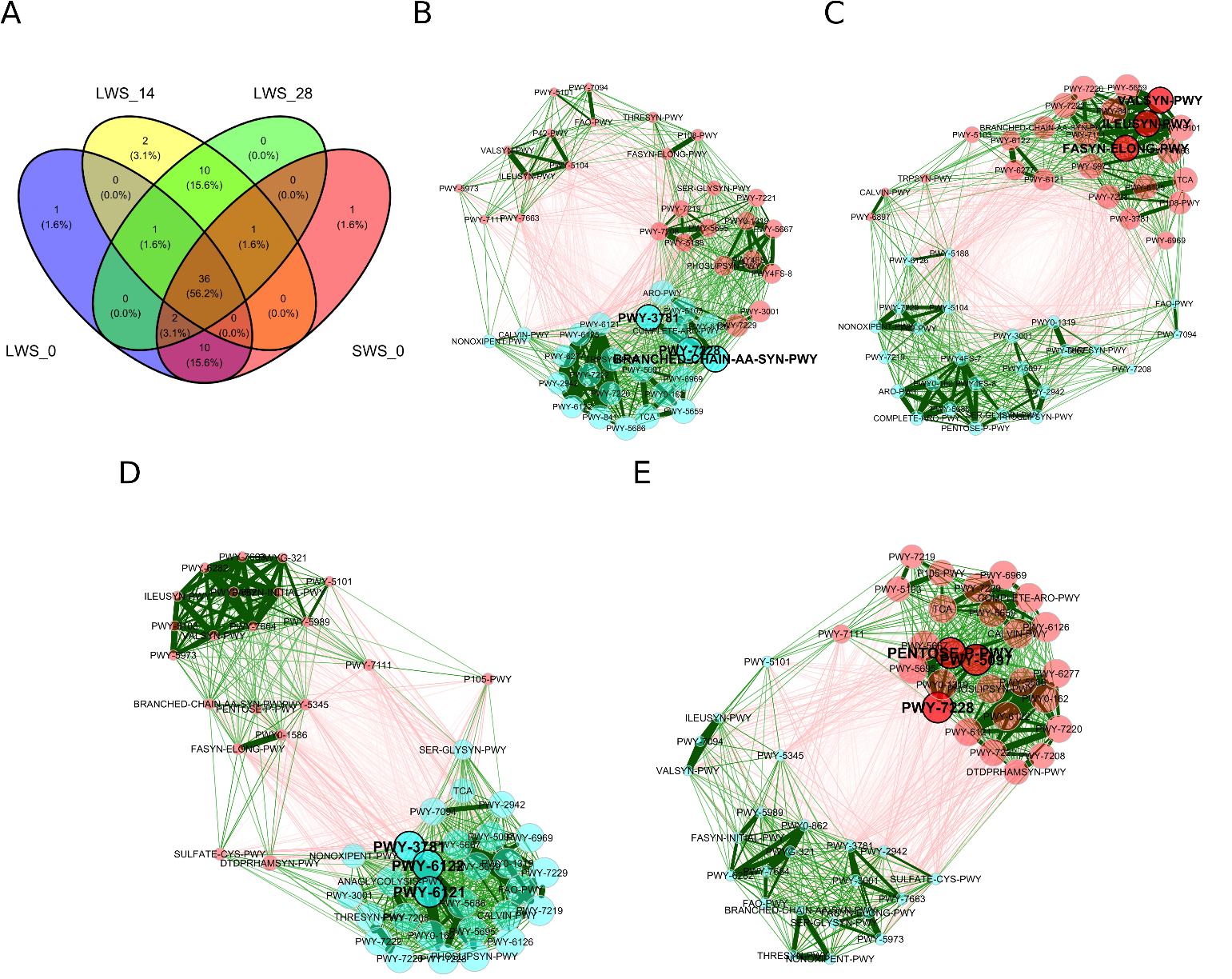

**Supplementary Figure S19.** **Network analysis of the top 50 most annotated functions for the WS environmental soil sample**. A. Venn diagram showing the overlap of the top 50 most annotated functions for each condition from the starting solid soil (SWS_0), the initial liquid NS-SESOM extraction at day 0 (LWS_0), and the NS-SESOM after 14 days (LWS_14) and 28 days (LWS_28) in continuous culture. B. Pathway-level network from the starting solid soil (SWS_0). C. Pathway-level NS-SESOM network for the initial liquid at day 0 (LWS_0). (D) Pathway-level NS-SESOM network after 14 days. (LWS_14). (E) Pathway-level NS-SESOM network after 28 days. (LWS_28). Red edges represent negative associations while green edges represent positive associations. Node colors represent clusters. Node size is scaled by eigenvector centrality.

Supplementary Figure S20.

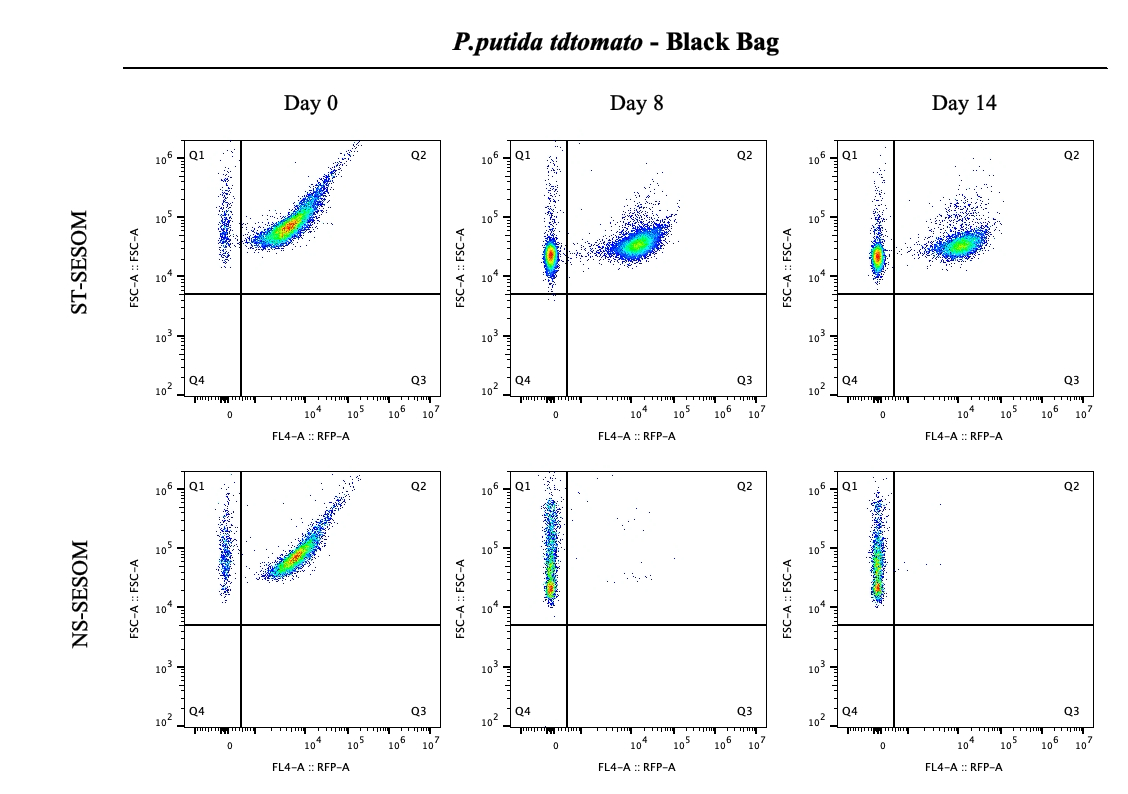

**Supplementary Figure S20. Flow cytometry dot plots and quadrant gating strategy for detection of *Pseudomonas putida* tdTomato in Black Bag ST-SESOM and NS-SESOM samples across timepoints.**Flow cytometry dot plots showing the gating strategy used to identify *P. putida* tdTomato cells in Black Bag ST-SESOM and NS-SESOM samples at Day 0, Day 8 and Day 14. A quadrant gate was applied using RFP-A fluorescence intensity on the x-axis and forward scatter (FSC-A) on the y-axis. Quadrant Q1 represents non-fluorescent events (RFP-negative), while Q2 contains RFP-positive *P. putida* tdTomato cells. This gating approach was used to distinguish bacterial populations and evaluate fluorescence-based detection of *P. putida* in each SESOM condition over time

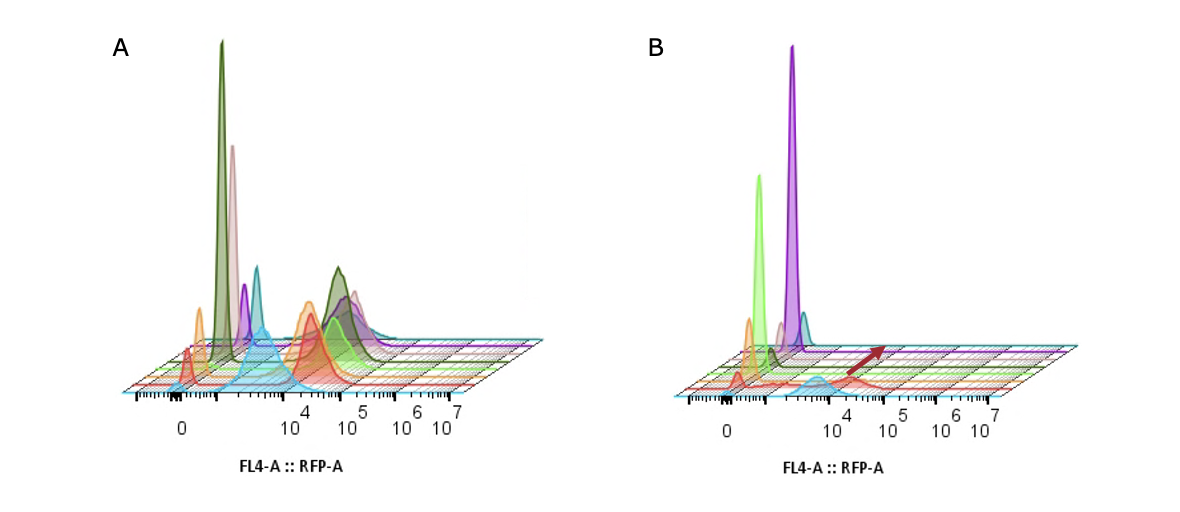

**Supplementary Figure S21. Flow cytometry fluorescence histograms of Pseudomonas putida tdTomato in Black Bag ST-SESOM and NS-SESOM over time.** Fluorescence intensity histograms showing tdTomato signal distribution of P. putida cells in (A) ST-SESOM and (B) NS-SESOM extracted from Black Bag soil. Histograms include samples collected every other day from Day 0 to Day 14 and are overlaid to visualize temporal changes in fluorescence. In panel A, tdTomato-positive populations consistently display a distinct peak around 10⁴ fluorescence units across all timepoints, while a separate peak near zero represents non-fluorescent events. In panel B, a small tdTomato-positive peak is observed at Day 0 and Day 2, followed by a flattened fluorescence distribution from Day 4 onward, indicating a loss of detectable fluorescent events over time. The red arrow highlights the shift from early timepoints toward reduced fluorescence intensity in NS-SESOM samples.
